# Rapid *de novo* assembly of the European eel genome from nanopore sequencing reads

**DOI:** 10.1101/101907

**Authors:** Hans J. Jansen, Michael Liem, Susanne A. Jong-Raadsen, Sylvie Dufour, Finn-Arne Weltzien, William Swinkels, Alex Koelewijn, Arjan P. Palstra, Bernd Pelster, Herman P. Spaink, Guido E. van den Thillart, Ron P. Dirks, Christiaan V. Henkel

**Affiliations:** ZF-screens B.V., J.H. Oortweg 19, 2333 CH Leiden, The Netherlands; Institute of Biology, Leiden University, Sylviusweg 72, 2333 CC Leiden, The Netherlands; Muséum National d’Histoire Naturelle, Sorbonne Universités, Research Unit BOREA, Biology of Aquatic Organisms and Ecosystems, CNRS, IRD, UCN, UA, 75231 Paris Cedex 05, France; Norwegian University of Life Sciences, Faculty of Veterinary Medicine, Department of Basic Science and Aquatic Medicine, PO Box 8146 Dep, 0033 Oslo, Norway.; DUPAN, PO Box 249, 6700 AE Wageningen, The Netherlands.; Animal Breeding and Genomics Centre, Wageningen Livestock Research, Wageningen University & Research, De Elst 1, 6708 WD Wageningen, The Netherlands.; Institute of Zoology and Center for Molecular Biosciences, University of Innsbruck, Innsbruck, Austria.; University of Applied Sciences Leiden, Zernikedreef 11, 2333 CK Leiden, The Netherlands.; Generade Centre of Expertise in Genomics, PO Box 382, 2300 AJ Leiden, The Netherlands.

**Keywords:** nanopore sequencing, genome assembly, eels, TULIP

## Abstract

We have sequenced the genome of the endangered European eel using the MinION by Oxford Nanopore, and assembled these data using a novel algorithm specifically designed for large eukaryotic genomes. For this 860 Mbp genome, the entire computational process takes two days on a single CPU. The resulting genome assembly significantly improves on a previous draft based on short reads only, both in terms of contiguity (N50 1.2 Mbp) and structural quality. This combination of affordable nanopore sequencing and light-weight assembly promises to make high-quality genomic resources accessible for many non-model plants and animals.

## Background

Just ten years ago, having one’s genome sequenced was the privilege of a handful of humans and model organisms. Spectacular improvements in high-throughput technology have since made personal genome sequencing a reality and prokaryotic genome sequencing routine. In addition, sequencing the larger genomes of non-model eukaryotes has opened up a wealth of information for plant and animal breeding, conservation, and fundamental research.

As an example, we and others [1–3] have previously established genomic resources for the European eel (*Anguilla anguilla*), an iconic yet endangered fish species that remains resistant to efficient farming in aquaculture [4, 5]. A draft genome [2], several transcriptomes (e.g. [1, 3, 6–10]), and reduced representation genome sequencing [11] have already shed light on its evolution and developmental biology [2, 12, 13], endocrinological control of maturation [7, 8], metabolism [14], disease mechanisms [10], and population structure [15, 16], thereby supporting both breeding and conservation efforts. However, compared to established model organisms, funds for eel genomics are naturally limited, and consequently the quality of current genome assemblies of *Anguilla* species is modest at best by today’s standards (Table 1).

**Table 1.**
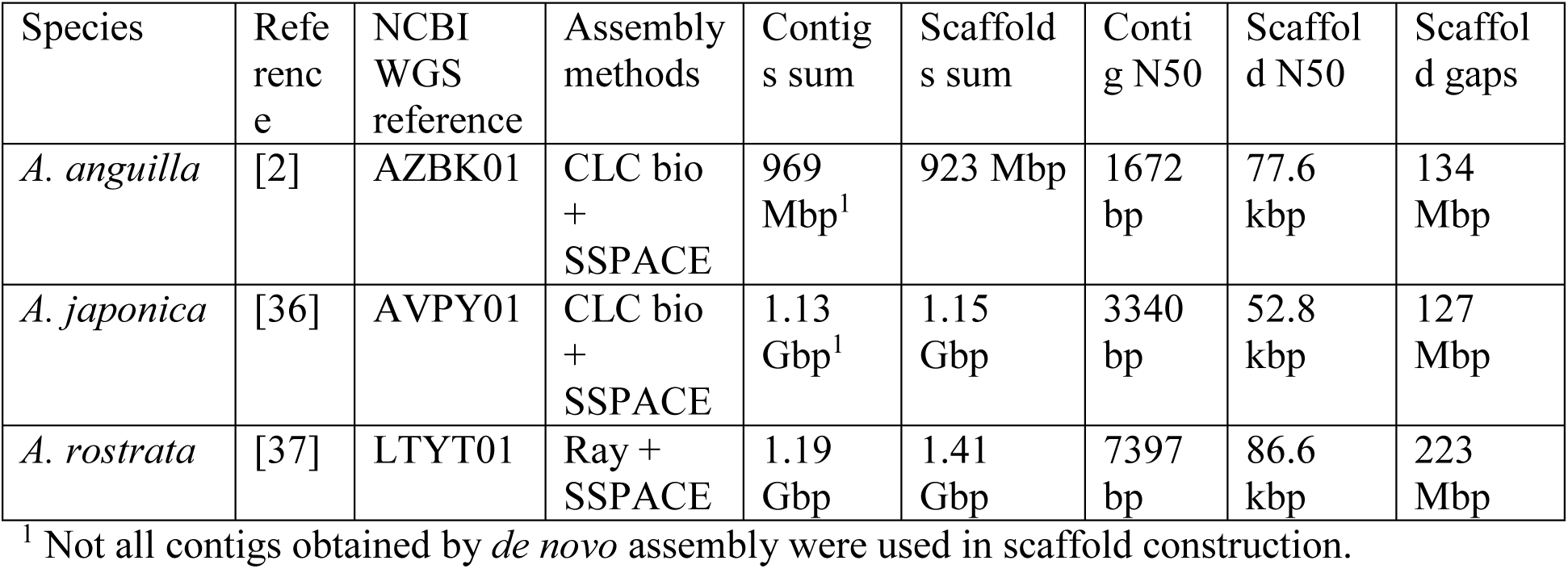
Previous genome assemblies of *Anguilla* species

The recent availability of affordable long-read sequencing technology by Oxford Nanopore Technologies (ONT, [17]) presents excellent opportunities for generating high-quality genome assemblies for any organism (for examples, see [18]). Flowcells for the miniature MinION sequencing device employ a maximum of 512 nanopores concurrently for reading single-stranded DNA at up to 450 nucleotides per second, resulting in several gigabases of sequence during a two day run. As the technology does not rely on PCR or discrete strand synthesis events, DNA fragments can be of arbitrarily long length. The single-molecule reads are of increasingly good quality, with a sequence identity of ~75% for the older R7.3 chemistry [17], to ~89% for the newer R9 chemistry (MinION Analysis and Reference Consortium, in preparation). Optionally, DNA can be read twice (along both strands) to yield a consensus ‘2D’ read of higher accuracy (up to ~94% for R9).

In contrast to short reads, long reads offer the possibility to span repetitive or otherwise difficult regions in the genome, resulting in strongly reduced fragmentation of the assemblies. This potential advantage does require the deployment of dedicated genome assembly algorithms that are aware of long-read characteristics. In addition, as single-molecule long-read technologies (by both PacBio and ONT) do suffer from reduced sequence identity, this likewise needs to be addressed by post-sequencing bioinformatics [19–21]. Dealing with these challenges has reinvigorated research into genome assembly methodology, resulting in several novel strategies [22–26].

However, when dealing with large eukaryotic genomes, the computational demands for long-read assembly are often higher than for short reads (using De Bruijn-graphs), even though the raw data are more informative of genome structure. Especially now that sequencing very large plant and animal genomes is finally becoming both technologically feasible and affordable, the computational costs may turn out to be prohibitive. For example, using the state-of-the-art Canu assembler [23], assembling a human genome from long PacBio reads takes thousands of CPU hours, or several days on a computer cluster. As scaling behavior is approximately quadratic with genome size, assembling a salamander [27] or lungfish [28] genome dozens of gigabases long would require several years on a cluster.

We are currently developing a computational pipeline specifically intended for future sequencing of extremely large tulip genomes (up to 35 Gbp, [29]). Here, we use a prototype of this algorithm to assemble a new version of the European eel genome, based on Oxford Nanopore sequencing. This entire computational process takes two days on a desktop computer, and yields an assembly that is two orders of magnitude less fragmented than the previous Illumina-based draft.

## Results

### Eel genome sizes and previous assemblies

Before launching a genome sequencing effort, an estimate of the size of the genome of interest is needed. For the genus *Anguilla*, several studies have used flow cytometry and other methods to arrive at C-values ranging from 1.01 to 1.67 pg [30], corresponding to haploid genome sizes in the 1–1.6 Gbp range for both *A. anguilla* and *A. rostrata*. We previously estimated a genome size of approximately 1 Gbp for *A. anguilla*, using human cells as a reference [2].

Based on their assembled genomes, *Anguilla* species exhibit a similarly wide range of apparent genome sizes (see Table 1). These draft assemblies are all based on previous-generation short-read technology, and relied on Illumina mate pairs to supply long-range information used in scaffolding. The resulting assemblies remain highly fragmented, with low N50 values even considering the technology used.

We therefore examined *k*-mer profiles in the raw Illumina sequencing data, which can provide an estimate of the length of the haploid genome [31, 32]. Surprisingly, the predicted genome sizes are considerably – but consistently – smaller than previously estimated or assembled (Table 2 and Fig. S1). In addition, all three examined genomes contain high levels of heterozygosity.

**Table 2.** *Anguilla* genome size predictions

| Species | Haploid genome size<sup>1</sup> | Repetitive fraction<sup>1</sup> | Heterozygous fraction<sup>1</sup> |
| --- | --- | --- | --- |
| *A. anguilla* | 854.0–866.5 Mbp | 15.5–20.0% | 1.48–1.59% |
| *A. japonica*<sup>2</sup> | 1.022 Gbp | 38.7% | 2.74% |
| *A. rostrata* | 799.0–813.0 Mbp | 12.2–16.9% | 1.50–1.60% |

<sup>1</sup> Ranges are the minimum and maximum values reported for three model fits at different $k$ -mer lengths. Apparent repetitive sequence decreases with $k$ -mer length, and heterozygosity increases with $k$ -mer length.
<sup>2</sup> For *A. japonica*, the model did not converge in most cases, presumably because of low coverage. These results are for $k = 19$ .

### Nanopore sequencing

We isolated DNA for long-read sequencing from the blood and liver of a fresh female European eel. Using three different generations of the ONT chemistry for the MinION sequencer, we generated 15.6 Gbp of raw shotgun genome sequencing data (see Table 3 and Fig. 1). Assuming an 860 Mbp haploid size, this corresponds to approximately 18-fold coverage of the genome. The bulk of the sequence is in long or very long reads (up to hundreds of thousands of nucleotides), although a fraction is composed of very short reads or artifacts (e.g. 6 bp reads, Fig. 1). We used all raw reads for subsequent genome assembly.

**Table 3.**
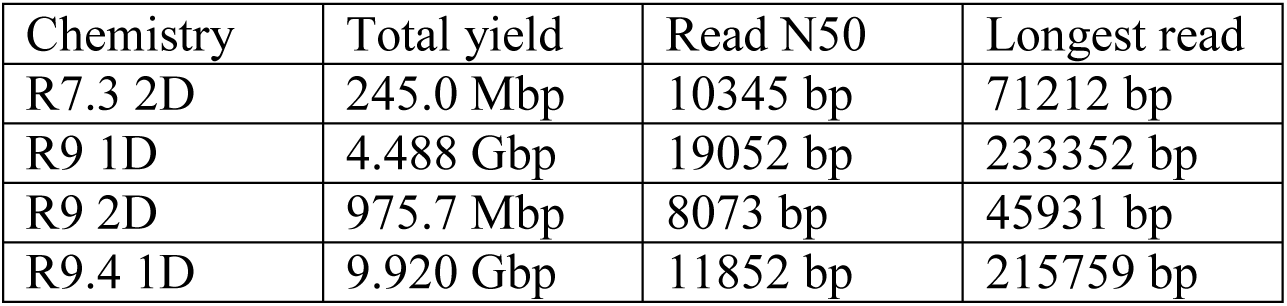
Nanopore sequencing

| Chemistry | Total yield | Read N50 | Longest read |
| --- | --- | --- | --- |
| R7.3 2D | 245.0 Mbp | 10345 bp | 71212 bp |
| R9 1D | 4.488 Gbp | 19052 bp | 233352 bp |
| R9 2D | 975.7 Mbp | 8073 bp | 45931 bp |
| R9.4 1D | 9.920 Gbp | 11852 bp | 215759 bp |

**Fig. 1.**
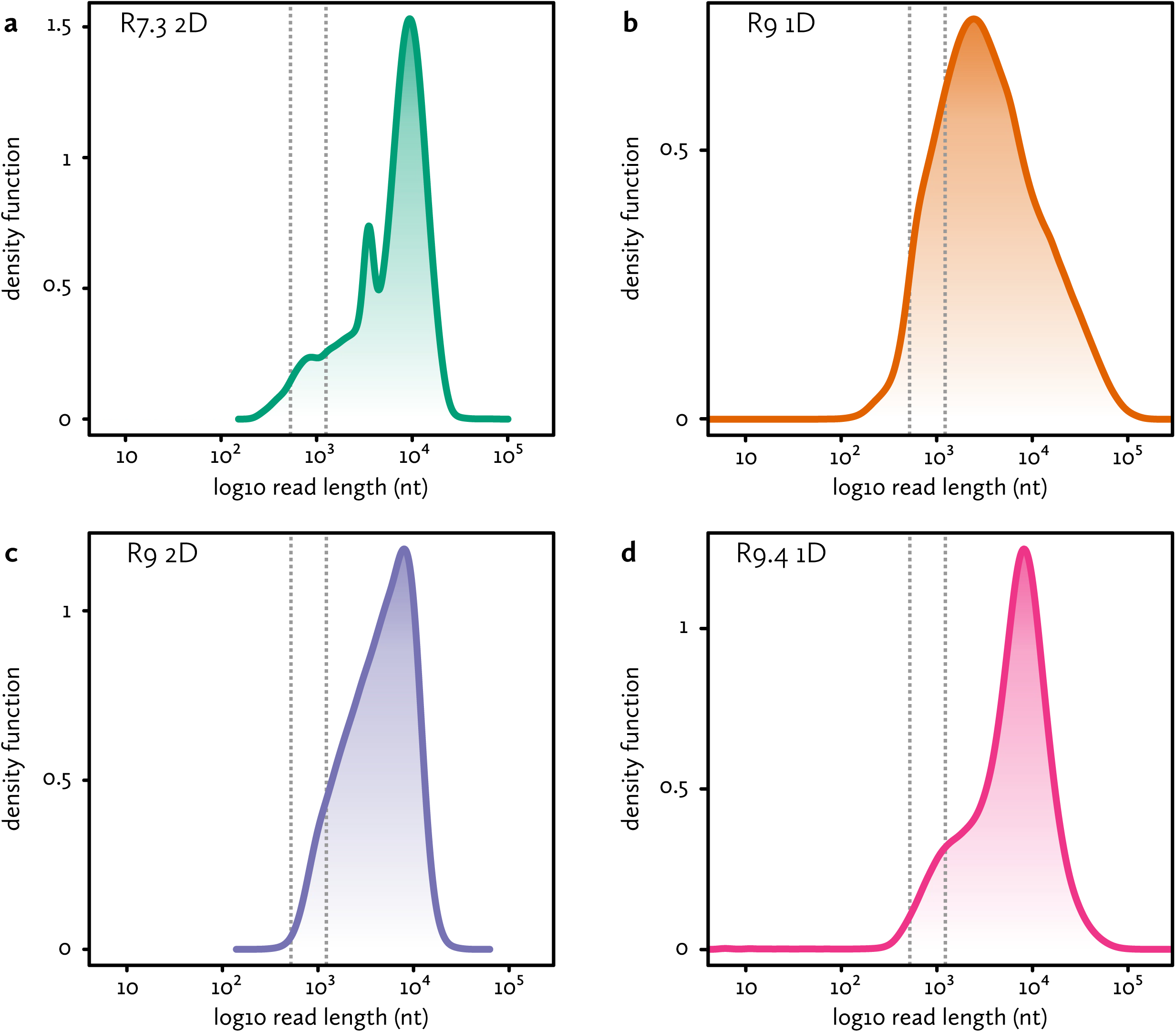
Nanopore sequencing Shown are the sequenced fragment size distributions for the **a** R7.3 chemistry 2D reads, **b** R9 chemistry 1D reads, **c** R9 chemistry 2D reads and **d** R9.4 chemistry 1D reads. Dotted lines indicate the minimum (542 bp) and typical (1270 bp) read lengths that can be used for linking two seeds in the 0.29× coverage 285 bp set. The minimum length is 2×285 bp with no more than 10% overlap between seeds. The typical length assumes an average of one seed per 985 bp (genome size divided by number of seeds).

### Assembly strategy

We assembled the long nanopore sequencing reads using a prototype of an assembly strategy we are developing for very large genomes (M. Liem and C. Henkel, in preparation), named TULIP (for *The Uncorrected Long-read Integration Process*). Briefly, it takes two shortcuts compared to the hierarchical approach [20–24]. First of all, like Miniasm [25], TULIP does not correct noisy single-molecule reads prior to assembly, but relies on a discrete post-assembly consensus correction application, e.g. Racon [19] or Pilon [33, 34]. Secondly, it does not perform an all-versus-all alignment of reads, but instead aligns reads to a sparse reference (of ‘seed’ sequences) that is representative for the genome.

Fig. 2a illustrates the steps we have taken to assemble the European eel genome. In this case, we employed previously generated Illumina shotgun sequencing reads as sparse seeds. Using a *k*-mer counting table, we identified merged read pairs that are suitably unique in the genome. Using strict criteria (see Methods), we could select 5019778 fragments of 270 bp, or 873058 of 285 bp, corresponding to 1.58-fold or 0.29-fold coverage of the genome, respectively. We subsequently used several random subsets of these fragments as a reference to align long nanopore reads against.

Using a custom script, we constructed a graph based on these alignments, in which the seed sequences are nodes, and edges represent long read fragments (Fig. 2b). A connection between two seeds indicates they co-align to a long read, and are therefore presumably located in close proximity in the genome. In theory, perfect alignments of very long reads to unique seeds should organize both sets of data into linear scaffolds.

However, because of the errors still present in long nanopore reads, the alignments are imperfect, with missed seed alignments making up the bulk of ambiguities in the seed graph (i.e. forks and joins in the seed path). Additional uncertainties are introduced by spurious alignments and residual apparently repetitive seeds. The tangles these cause in the graph can be recognized locally, and are removed during a graph simplification stage (Fig. 2c). TULIP will visit every seed that has multiple in- or outgoing connections, and attempt to simplify the local graph topology by removing connections. For example, if a single seeds fails to align to a single nanopore read, this will introduce a ‘triangle’ in the graph (Fig. 2c, top example), in which the neighbouring seeds now share a direct connection (based on that single read). If the intermediate seed fits between the neighbouring seeds, TULIP will then remove the connection spanning the intermediate seed. If after this stage a seed still has too many connections, it might represent repetitive content and its links are severed altogether (Fig. 2c, second example).

**Fig. 2.**
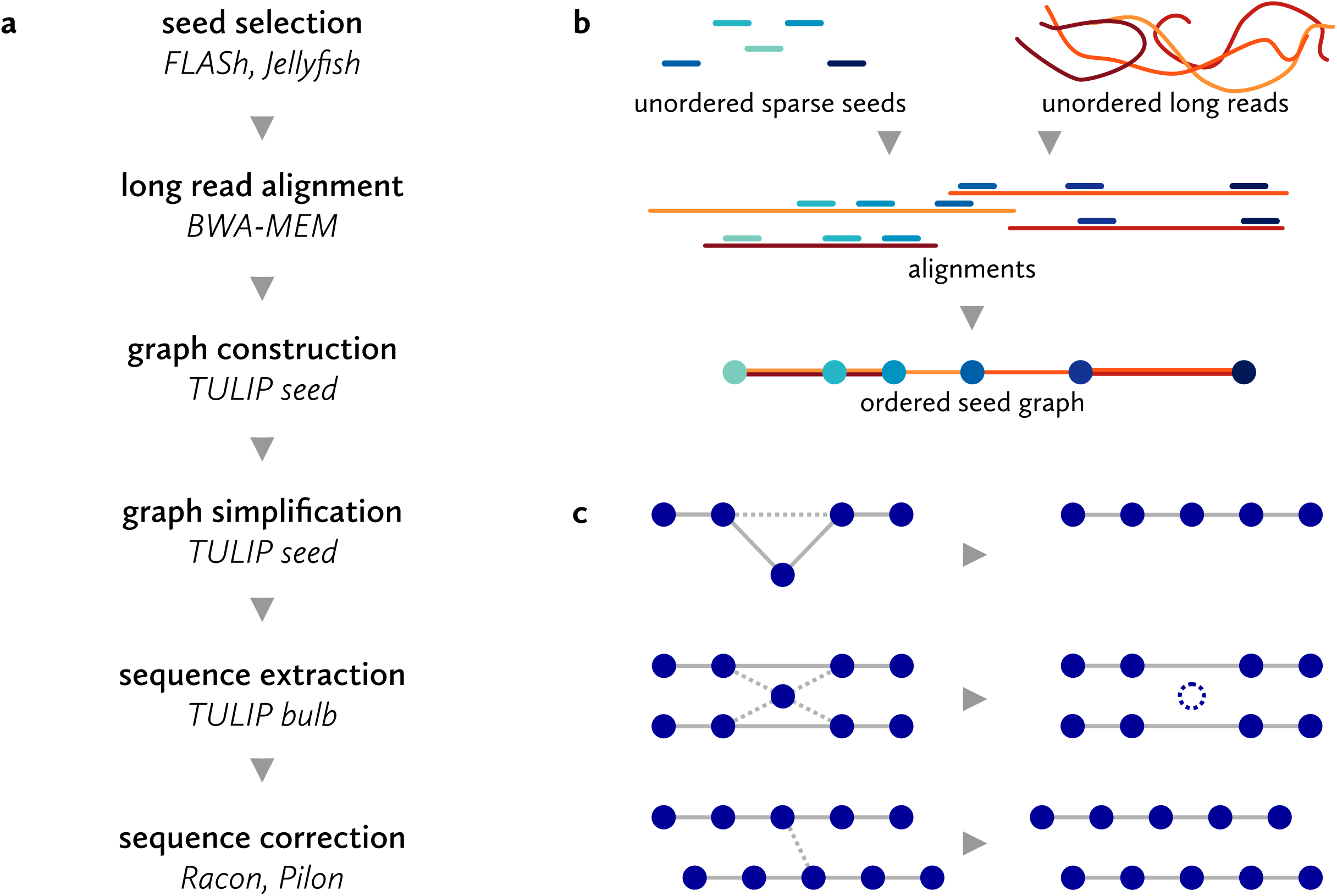
Assembly strategy **a** Stages in TULIP. **b** Graph construction based on long read alignments to short seeds. Seeds are included in the graph as nodes if they align adjacent to each other to a long read. The apparent distance between the seeds is included as an edge property, as is the amount of evidence (i.e. number of alignments supporting the connection). **c** The initial seed graph based on alignments contains ambiguities, caused by missed alignments, repetitive seed sequences and spurious alignments. These are removed during the initial layout process, resulting in linear scaffolds. Where possible, these scaffolds are subsequently linked by further unambiguous long-distance co-alignments to long reads.

Finally, unambiguous linear arrangements of seeds can be extracted from the graph. Fig. 3 illustrates a small fragment of the actual seed graph, with final linear paths (scaffolds) and removed connections indicated.

**Fig. 3.**
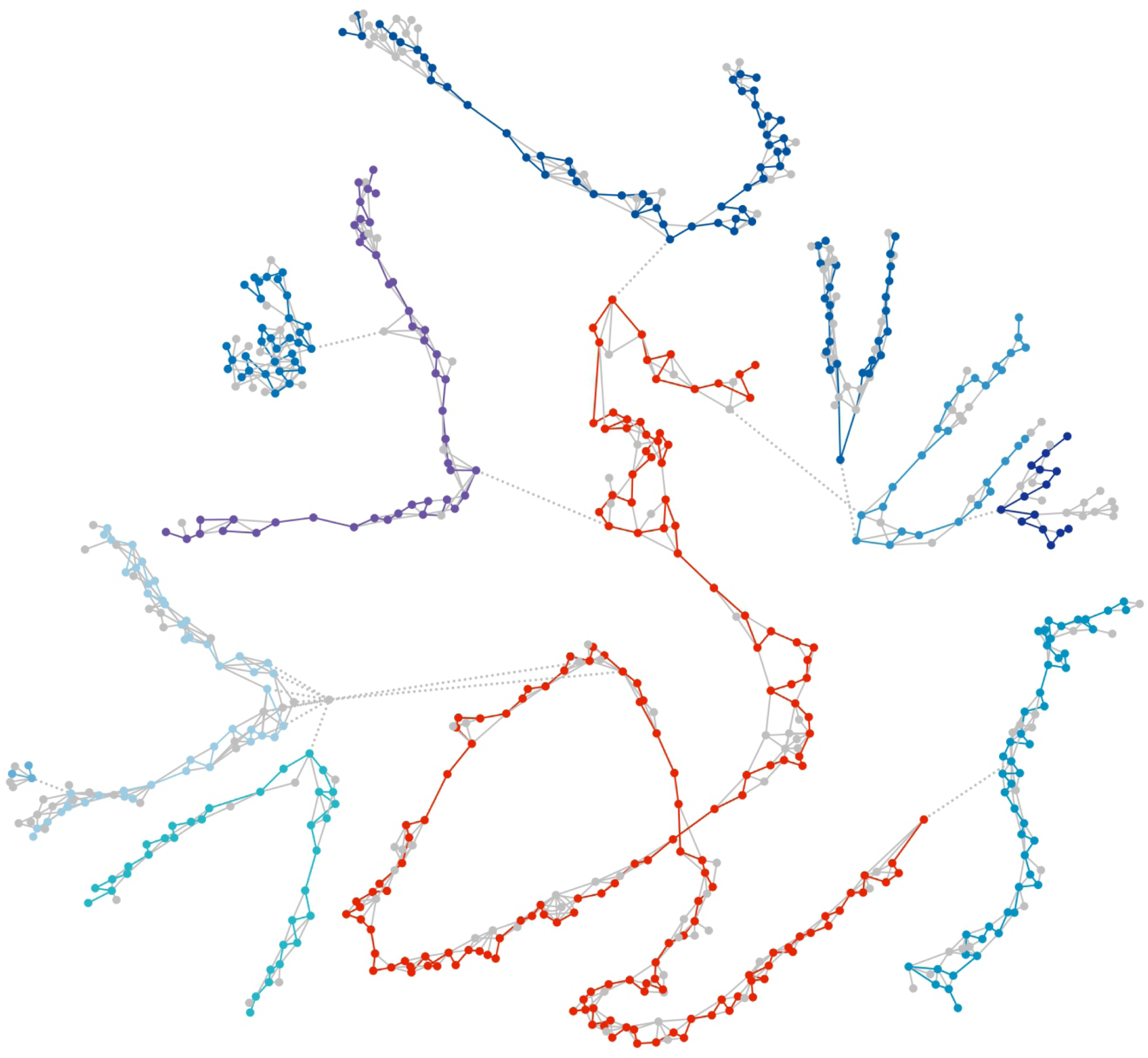
Scaffolds were extracted from a graph consisting of seed sequences (nodes) linked by nanopore reads (edges). Here, a small final scaffold (number 2231, 252.2 kbp) is shown in red in the context of the initial seed graph (all seeds at a distance of up to ten links from the final scaffold). Fragments of ten other scaffolds (blues) are directly or indirectly connected to scaffold 2231 by a few incorrect links (dotted lines). Seeds and links removed during graph simplification are shown in grey. Scaffolds can be discontinuous in the initial graph, as additional long-distance links are added in a later stage. The graph was visualized using Cytoscape (version 3.4.0).

These ordered seed scaffolds do not yet contain sequence data. These can subsequently be added from the original nanopore reads and alignments, resulting in uncorrected scaffold sequences. The scaffolds are exported bundled with their constituent nanopore reads, and can be subjected to standard nanopore sequence correction procedures.

### Assembly characteristics

We used several combinations of short seed sequences and aligned nanopore reads to optimize the assembly process. In most cases, we did not complete the entire assembly process by adding actual nanopore sequence. Therefore, distances between seeds (and scaffold lengths) are means based on multiple nanopore reads. Adding specific sequence (and subsequently correcting scaffolds) can change these figures slightly. Table 4 lists the assembly statistics for these experimental runs.

**Table 4.** *A. anguilla* genome assemblies using TULIP

| Seed size | Seed number | Read selection | Scaffold N50<sup>1</sup> | Nr. of scaffolds | Assembly size<sup>1</sup> |
| --- | --- | --- | --- | --- | --- |
| 285 bp | 873k | 100% | 1170852 bp | 2366 | 849.7 Mbp |
| 285 bp | 873k | 75% | 697683 bp | 3531 | 839.0 Mbp |
| 285 bp | 873k | 50% | 341223 bp | 6919 | 815.0 Mbp |
| 285 bp | 873k | 25% | 90534 bp | 21764 | 730.4 Mbp |
| 285 bp | 437k | 100% | 719956 bp | 3173 | 802.6 Mbp |
| 285 bp | 218k | 100% | 361910 bp | 4889 | 709.6 Mbp |
| 270 bp | 1746k | 100% | 1185122 bp | 2805 | 875.6 Mbp |
| 270 bp | 1310k | 100% | 1300479 bp | 2317 | 866.7 Mbp |
| 270 bp | 873k | 100% | 1176872 bp | 2330 | 851.0 Mbp |
| 270 bp | 437k | 100% | 711245 bp | 3132 | 802.6 Mbp |

<sup>1</sup> Sizes based on mean distances between seeds.

Both the contiguity and size of the assembly clearly improve upon adding more nanopore data (Fig. 4a, b). This suggests that at 18-fold coverage of this genome, and using the particular blend of data types available here, the assembly process is still limited by the total quantity of long read data.

For the seeds, we investigated the effects of seed length (270 or 285 bp), as well as seed density (fractions and multiples based on the 873058 fragments available at 285 bp). There does not appear to be a clear advantage to choosing either 270 or 285 bp seeds. At identical densities, the two possibilities yield comparable assemblies in terms of size and contiguity. For seed density, there does appears to be an optimum. As expected, low densities result in fragmentation and incompleteness (Fig. 4c, d). The assemblies with the highest seed density (1.3 or 1.7 million 270 bp sequences) do yield the highest N50 and assembly sum (Table 4), but also exhibit increased fragmentation compared to lower seed densities. As Fig. 4c shows, the main difference with those assemblies is the appearance of many small scaffolds at high seed numbers.

Accidentally, in this case the optimal seed density is around the ‘full’ set of 873058 fragments, of either 270 or 285 bp. Both also yield an assembly that is close to the estimated genome length. We selected the 285 bp version as a candidate for an updated reference genome for the European eel.

Fig. 4 summarizes several characteristics of the candidate assembly (before sequence addition or correction). The length distribution of the 2366 scaffolds (Fig. 4a) shows they range in size between 431 bp and 8.7 Mbp. The lower boundary is expected, as a minimal scaffold has to consist of at least two 285 bp seeds, and the graph construction was executed with parameters allowing limited overlap between seeds. The cumulative scaffold length distributions (Fig. 4b) show that a considerable fraction of the genome is included in large scaffolds, with 232 scaffolds larger than a megabase constituting 56% of the assembly length. Seeds in the final scaffolds are connected by on average 7.4 nanopore read alignments. As can be seen in Fig. 4e, links removed during the graph simplification stage (mostly based on local graph topology only) were predominantly those supported by less evidence.

The final assembly retains 637792 seeds of 285 bp, equivalent to a maximum of 181.8 Mbp of Illumina-derived sequence. If the seed distribution is assumed to be essentially random (with local genomic architecture responsible for exceptions), the initial 873058 seeds should be spaced at a mean interval of 700 bp. As seeds are removed during simplification, larger ‘gaps’ filled with nanopore-derived sequence should appear. However, as Fig. 4f shows, gap lengths are heavily biased towards low and negative lengths (i.e. overlapping seeds). In this case, this could be an artifact of the very stringent seed selection procedure.

**Fig. 4.**
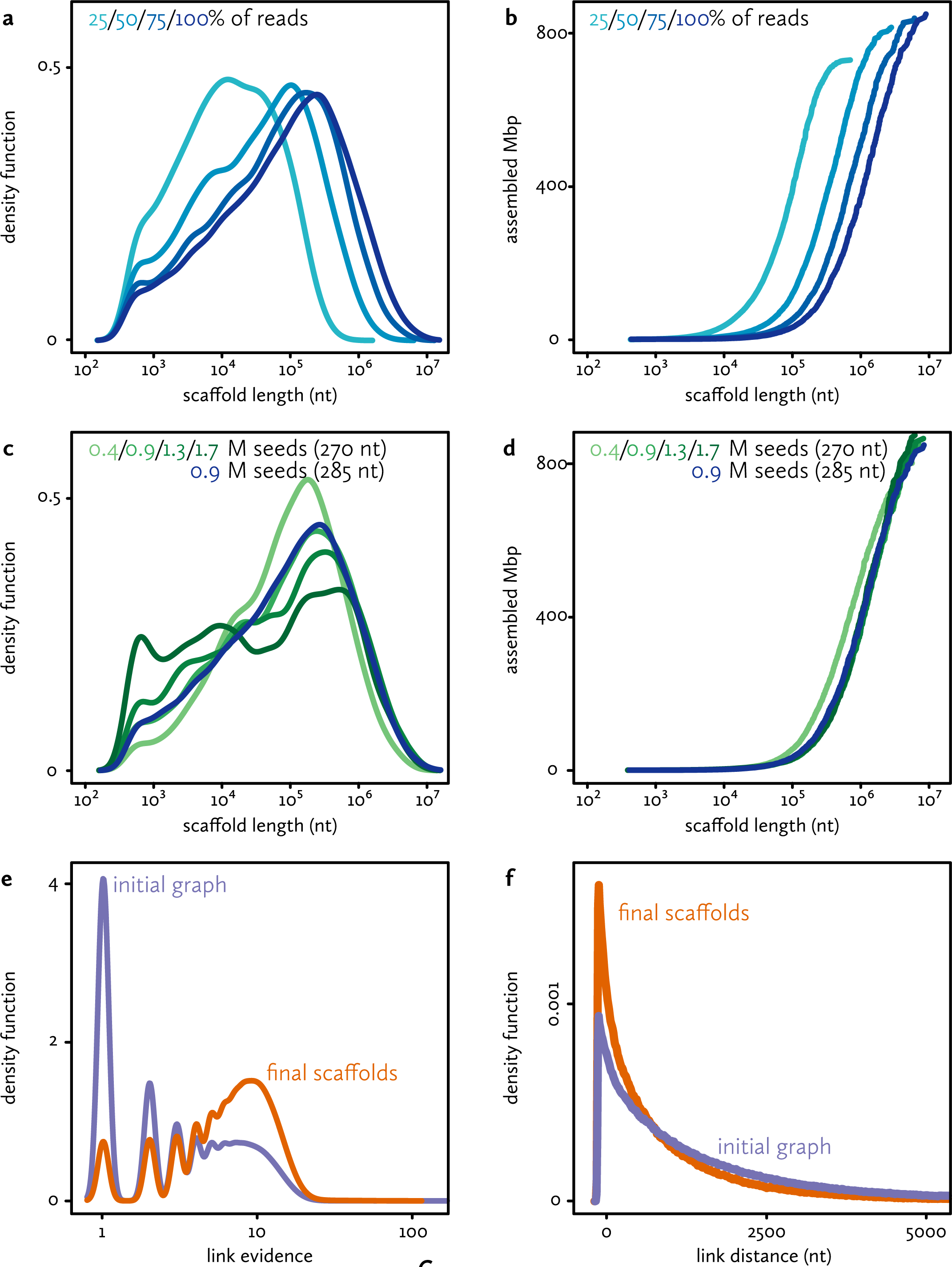
Characteristics of the final assembly **a** Size distribution of final scaffolds, based on 285 bp seeds. Colours indicate alternative assembly runs, using subsets of the long read data. **b** Cumulative size of the final scaffolds, sorted by size. **c** and **d** Size distributions and cumulative size distributions for final scaffolds, based on both 270 and 285 bp seeds. Colours indicate alternative assembly runs, using different seeds sets. **e** Link evidence distribution in the initial graph (purple) and the final graph (orange) for the candidate assembly (285 bp seeds). **f** Distances between seeds in the initial graph (purple) and the final graph (orange) for the candidate assembly (285 bp seeds).

### Assembly quality

In order to assess its completeness and structural correctness, we added nanopore sequence to the selected TULIP assembly and aligned it to the Illumina-based draft genome [2]. As a high-quality reference genome for the European eel is not yet available, such a comparison need take into account the possibility of error in either assembly. However, with appropriate caution, agreement between the assemblies – which are completely independent in both sequencing data and assembly algorithms – can confirm the integrity of both.

Fig. 5a shows a full-genome alignment of the new (uncorrected) nanopore-based assembly to the 2012 draft [2], based on best pairwise matches. This confirms that at this large scale, all sequence in the new assembly is also present in the older assembly. At first sight, the converse does not appear to be the case: the Illumina-based draft is 923 Mbp in size, and contains approximately 96 Mbp in scaffolds that have no reciprocal best match in the nanopore assembly (863.3 Mbp after sequence addition, see Table 5). However, the non-matching sequences consist almost exclusively of very small scaffolds (mean/N50 664/987 bp). Since the Illumina-based draft assembly also contains 134 Mbp in gaps, these small scaffolds are plausibly sequences that could not be integrated correctly during the SSPACE scaffolding process [35, 36]. Both assemblies therefore roughly span the entire predicted genome of 860 Mbp.

**Table 5.** Characteristics of the *A. anguilla* candidate assembly

| Statistic | Value | Note |
| --- | --- | --- |
| Number of scaffolds | 2366 |  |
| Seed graph scaffold N50 | 1.17 Mbp | cf. Table 4 |
| Seed graph assembly sum | 849.7 Mbp | cf. Table 4 |
| Uncorrected scaffold N50 | 1.19 Mbp |  |
| Uncorrected scaffold sum | 863.3 Mbp |  |
| Racon scaffold N50 | 1.21 Mbp |  |
| Racon assembly sum | 881.3 Mbp |  |
| Pilon scaffold N50 | 1.23 Mbp |  |
| Pilon assembly sum | 891.7 Mbp |  |
| Alignment time | 7 hours | 1 thread<sup>1</sup> |
| Seed graph time | 51 minutes | 1 thread |
| Sequence addition time | 14 minutes | 1 thread |
| Racon correction time | ~22 hours | 1 thread<sup>1</sup> |
| Pilon correction time | ~24 hours | 1 thread<sup>1</sup> |

<sup>1</sup> These stages can be sped up by multithreading. For example, the actual alignment was run with four concurrent threads in 2 hours, 34 minutes.

Fig. 5b–f show detailed alignments, based on the 5 largest nanopore scaffolds (6.1–8.9 Mbp uncorrected) and their best matches only. These alignments confirm that in this sample both assemblies are mostly collinear, with the smaller Illumina draft scaffolds usually aligning end-to-end on the larger TULIP scaffolds. Therefore, both presumably reflect the actual genomic organization. However, at this level of detail several structural incongruities between both assemblies also become apparent (indicated by arrowheads). For 16 scaffolds from the 2012 draft, only part of the sequence is present in the selected TULIP scaffolds. In other words, at these loci both assembly protocols made different choices, based on the available sequencing information.

**Fig. 5.**
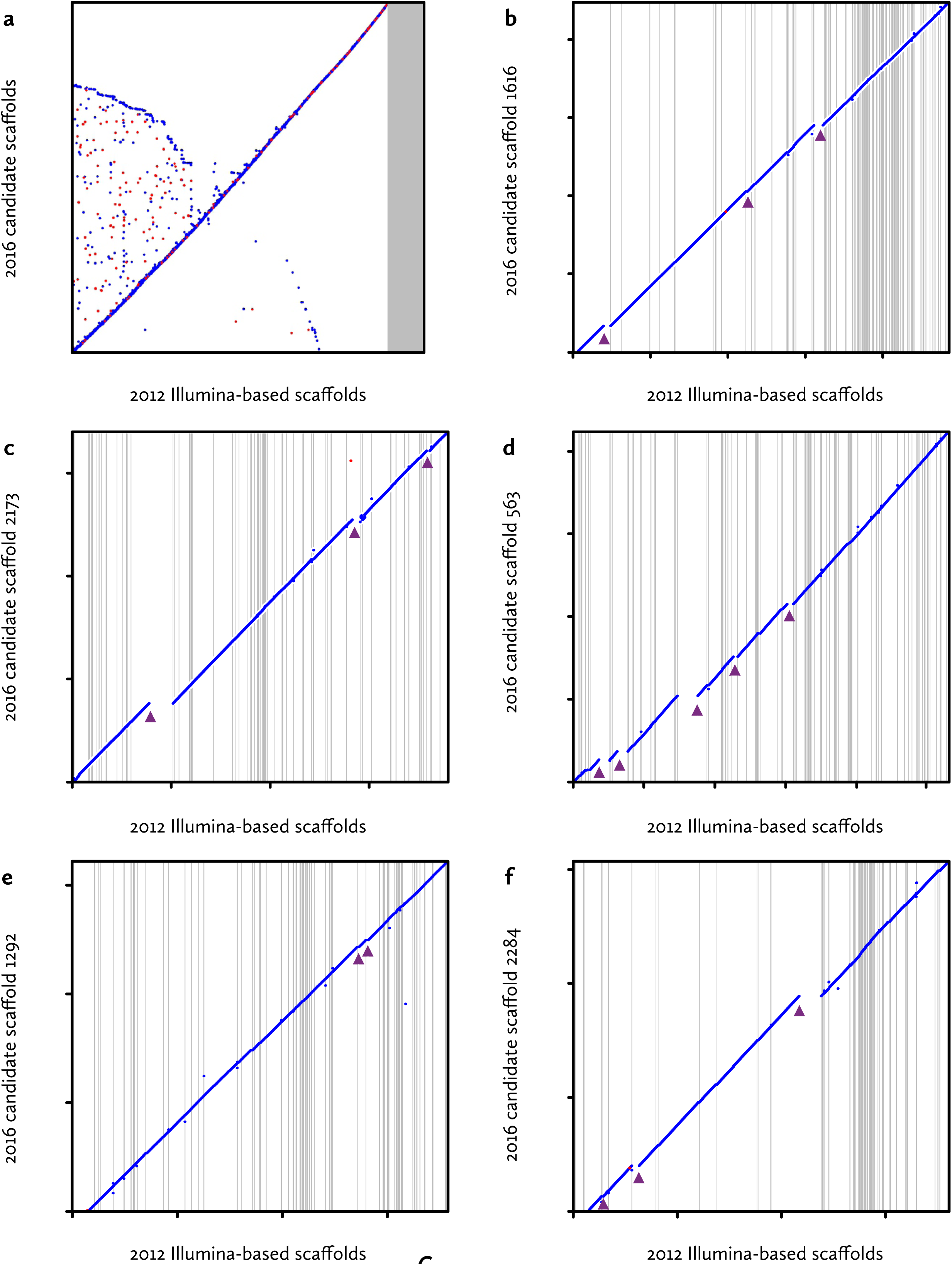
Full-genome alignment of the final assembly **a** The final uncorrected scaffolds (N50 = 1.19 Mbp, y-axis) were aligned to the 2012 *A. anguilla* assembly (N50 = 77.6 kbp, x-axis) using nucmer [51] with minimum match length 100, filtered for best pairwise matches between scaffolds (delta-filter −1), and plotted using the mummerplot --layout option. The grey area corresponds to small scaffolds in the 2012 assembly that are not part of a best reciprocal match. (**b**–**f**) More detailed alignments between the five largest nanopore scaffolds (y-axes) and their best matches in the 2012 draft assembly (x-axes). Grey horizontal and vertical lines indicate scaffold boundaries. These figures were generated in R (version 3.3.1) based on mummerplot output. 2012 draft scaffolds with minimal contributions to the overall alignment were removed manually. Arrowheads indicate discrepancies between both assemblies.

We therefore examined the evidence for the decisions made by TULIP. For each discrepancy, we examined the local neighbourhoods in the initial nanopore-based seed graphs (as in Fig. 3). If a draft scaffold is correct, at the inconsistency there should be multiple alternatives for the TULIP algorithm to choose from (Fig. S2). As these subgraphs (Fig. S3–S7) show, there is no evidence in the nanopore data for the older draft structure for any of the 16 cases examined. On the contrary, most local graph neighbourhoods appear relatively simple and support unambiguous scaffolding paths. The links at these suspect junctions are supported by at least two (average six) independent nanopore reads, which reduces the likelihood of accidental connections (caused by e.g. chimaeric reads).

Fig. S4. Local graph neighbourhoods of scaffold inconsistencies. For each of the inconsistencies identified in Fig. 5b–f, the local neighbourhood in the initial seed graph is shown (similar to Fig. 3 and Supplementary Fig. 2c). Red and green nodes represent seeds that align to the truncated old scaffold and its non-truncated neighbour, respectively. Grey nodes do not align to these scaffolds (or at least, not locally), yellow nodes align partially to two scaffolds. The final extracted TULIP scaffold paths are indicated by blue arrows. As in the draft the ‘red’ scaffolds do not end at the joins to the ‘green’ scaffolds, an alternative path possibility of continuing with ‘red’ seeds would be expected at this point. In none of the cases examined does this appear to be the case.

Alternatively, the order of the draft scaffolds in the alignments already suggests which of the two assemblies is correct. If one of the 16 problematic scaffolds were to reflect the legitimate genome structure, this error in the new assembly would usually also affect the next aligning scaffold. However, in almost all cases, the neighbouring draft scaffold aligns end-to-end. This suggests that either the TULIP assembly intermittently features very large rearrangements that accidentally always end at draft scaffold boundaries, or that the draft scaffolds are occasionally misconstrued.

Finally, the distribution of draft scaffolds along the nanopore-based scaffolds reveal an interesting pattern. The distribution of draft scaffold length along the genome is clearly non-random, with some regions assembled into just a few large scaffolds, whereas other regions (often up to a Mbp in size) are highly fragmented into very small scaffolds. This indicates that using short-read technology, certain genomic features are intrinsically harder to assemble than using long reads.

### Sequence correction

Currently, the ONT platform does not yield reads of perfect sequence identity. Like with PacBio data, therefore, at some point in the assembly process the single-molecule-derived sequence needs to be corrected by extracting a consensus from multiple reads covering every genomic position. Here, we opted for a standalone post-assembly correction step with Racon, which extracts a consensus from nanopore reads [19]. As some positions in the assembly are based on a single nanopore reads (Fig. 4e), in this case this correction may not be sufficient. Therefore, we subsequently corrected with Pilon, which extracts a consensus based on alignment of Illumina reads to the noisy sequence [33, 34]).

To assess the changes made by these correction algorithms, we counted and compared the occurrence of 6-mers in the draft Illumina-based assembly, the uncorrected TULIP assembly, and after correction (Fig. 6). These frequencies reveal several expected patterns, specifically a slight underrepresentation of high CG content in Illumina-based sequence (draft and Pilon), and an underrepresentation of homopolymer sequence in nanopore-based sequence (TULIP and Racon) [17]. Overall, the correction steps bring the sequence similarity of the nanopore-based assembly closer to the Illumina-based draft, with the final corrected assembly having a high correlation to the draft (Fig. 6 lower left panel).

**Fig. 6.**
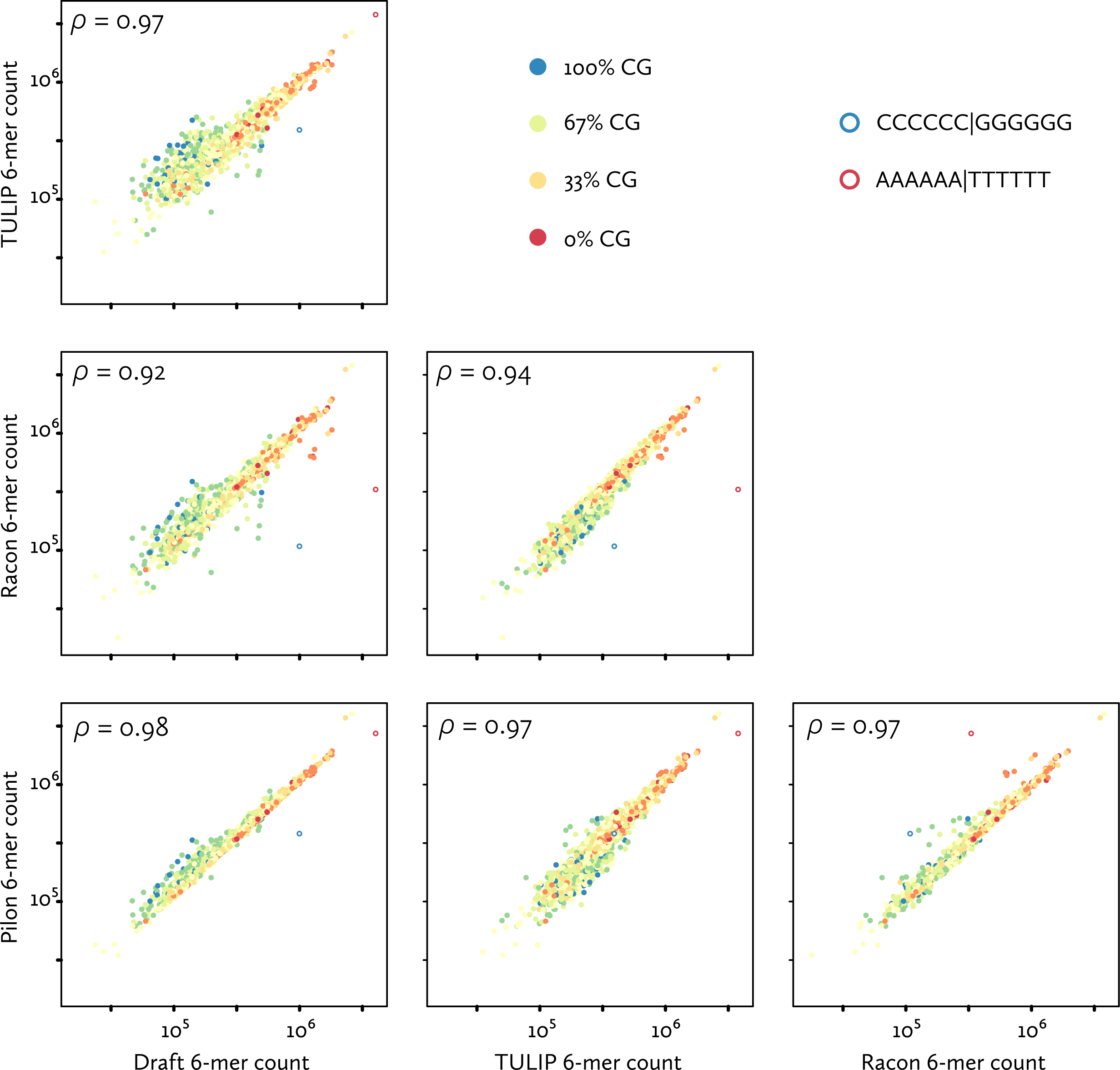
Sequence identity in nanopore-based assemblies The sequence similarity to the older draft of different stages of the nanopore assembly process (uncorrected TULIP, corrected by Racon, and additionally corrected by Pilon) is illustrated by 6-mer frequency counts (generated using Jellyfish). With every point a discrete 6-mer, colours indicate CG-content, and open circles indicate the two homo-6-mers. Scales are logarithmic. Also shown are Pearson correlation coefficients between the frequency distributions.

Sequence correction remains the most time-consuming stage of the assembly, requiring 22 and 24 hours (on a single CPU) for Racon and Pilon, respectively (Table 5). As TULIP bundles uncorrected scaffolds with its constituent nanopore reads, this process could still be sped up by parallelization, with individual scaffolds distributed over concurrent correction threads.

## Discussion

In this study, we have evaluated whether it is possible to sequence a vertebrate genome using nanopore long-read technology, and quickly assemble it using a relatively simple and lightweight procedure.

One of the most striking outcomes of this eel genome sequencing effort is the surprisingly close match between the genome size predicted from *k*-mer analysis (~860 Mbp) and the TULIP assembly (891.7 Mbp after corrections), and their distance from short-read-based assemblies. This can be explained either by the absence of a substantial fraction of the genome from the nanopore data or assembly, or by an artificially inflated genome size for the short-read assemblies. Full-genome alignment between both assemblies (Fig. 5a) suggests the latter phenomenon is at least partially responsible, as only tiny short-read scaffolds are absent from the long-read assembly.

An analysis of the short-read *A. anguilla* [2] and *A. japonica* [36] assembly procedures implies that the scaffolding process, based on mate pair data, is responsible for the introduction of numerous gaps (Table 1). In addition, at the time we discarded a considerable fraction of the initial contigs, which was composed primarily of very small contigs that appeared to be artefactual (based on low read coverage or very high similarity to other contigs). Plausibly, such contigs – and the high residual fragmentation of these assemblies – are the result of the high levels of heterozygosity in these genomes (Fig. S1).

Similar processes could also explain the even larger discrepancy between the predicted and assembled size of the recently published genome of the American eel *A. rostrata* (Table 1, [37]). As European and American eels interbreed in the wild [38], a large difference in genome size is unlikely – although it could also provide an explanation for the observed limited levels of gene flow between the species [16].

The whole-genome alignments between the Illumina draft and the new nanopore-based assembly (Fig. 5) also serve to confirm the structural accuracy of both. In a small sample (corresponding to of 4.2% of the genome), we observed 16 apparent assembly errors (Fig. 5b–f). In the absence of a high-quality reference, it is difficult to establish which assembly is correct. However, our analyses strongly suggest that in these cases the nanopore-based assembly is accurate. This is not unexpected: TULIP has access to far richer and more accurate sequencing information than SSPACE, which had to rely on 2×36 bp mate pair data. Under such circumstances, a low number of incorrect joins between contigs is inevitable [39]. In fact, considering the fact that the SSPACE scaffolds analyzed in Fig. 5b–f consist of on the order of ten thousand very small contigs, a result with only 16 errors signifies better scaffolding performance than expected [39].

In other aspects, the TULIP assembly is likely to be suboptimal. By design, scaffolds that could be merged based on long reads remain separate if these reads do not share a fortuitous seed alignment in the correct position. Similarly, large repetitive regions in the genome, as well as (sub)telomeric repeats will not always contain frequent 285 bp islands of unique sequence, and hence could be absent from the assembly. Although counterintuitive, this should not pose a major problem for some extremely large genomes. Survey sequencing indicates that the 32 Gbp axolotl genome contains mostly unique sequence [27], as do many tulip genomes (C. Henkel, unpublished data).

The selection of sparse seeds by the user adds an unusual level of flexibility to the assembly process. In an early phase of this study, we opted for essentially randomly placed Illumina-based seed sequences. This choice was motivated by their very high sequencing identity, which aids alignment quality when working with early, error-prone nanopore chemistries [17] However, with the speed at which the quality of reads produced by the ONT platform is improving [18], it should soon be possible to avoid such a hybrid assembly altogether. A natural choice for seed sequences would then be the ends of long reads.

Alternatively, seeds could be chosen to facilitate further sequence integration. If a high density genetic map is available for a species, map markers could serve as pre-ordered seeds. For example, with minor modifications, TULIP might be used to selectively add long read sequencing data only to single map marker bins (containing thousands of actual, unordered markers) resulting from a population sequencing strategy [40].

The bottleneck for such strategies lies in the interplay between marker density and nanopore read length, where the latter currently appears to be limited chiefly by DNA isolation protocols [41, 42]. Conceivably, in the near future, the problem of genome assembly from sequencing reads will all but disappear: abundant megabase-sized reads of high sequence identity are becoming conceivable, which should span the vast majority of recalcitrant regions in medium-sized genomes that remain a challenge to short- and medium-read technologies.

The fulfillment of such prophesies may still lie several years in the future. Therefore, we plan to further integrate and validate the candidate assembly generated here with long-range information obtained from optical mapping [43], in order to develop a high-quality reference genome for the troubled European eel.

## Conclusion

We have developed a new, simple methodology for the rapid assembly of large eukaryote genomes using a combination of long reads and short seed sequences. Using this method, we could assemble the 860 Mbp genome of the European eel using 18× nanopore coverage and sparse pre-selected Illumina reads in three hours on a modest desktop computer. Including subsequent sequence correction, the entire process takes two days. This yields an assembly that is essentially complete and of high structural quality.

## Methods

### Genome size estimation and k-mer analyses

We used Jellyfish version 2.2.6 [44] to count *k*-mers in sequencing reads and assemblies. In order to estimate genome size, we obtained frequency histograms for 19- to 25-mers in raw Illumina sequencing data. Reads were truncated to a uniform length of 76 nt, except for *A. japonica*, for which we used 100 nt (the model did not converge for short lengths). For the American eel, which has been sequenced at much higher coverage than the European and Japanese species, we used a subset of the available data (SRR2046741 and SRR2046672). Histograms were analyzed using the GenomeScope website [32] in order to obtain estimates for genome sizes, heterozygosity and duplication levels.

### Illumina seed selection

We selected unique seed sequences from 11.9 Gbp in sequence previously generated at 2×151 nt on an Illumina Hiseq 2000. Pairs were merged using FLASh [45], requiring a minimum of 15 nt terminal overlaps, resulting in 29.16% merged fragments. In these, 25-mers were counted using Jellyfish. We used a custom script to filter out all fragments that contained 25-mers occurring over 25 times in the remaining data. This corresponds to a maximum occurrence of approximately 6.25× in the 860 Mbp genome. Finally, fragments were selected based on size (either 270 nt or 285 nt).

### DNA purification

High MW chromosomal DNA was isolated from European eel blood and liver samples using a genomic tip 100 column according to the manufacturer’s instructions (Qiagen).

### MinION library preparation and sequencing

The genomic DNA was sequenced using nanopore sequencing technology. First the DNA was sequenced on R7.3 Flow Cells. Subsequently multiple R9 and R9.4 Flow Cells were used to sequence the DNA. For R7.3 sequencing runs we prepared the library using the SQK-MAP006 kit from Oxford Nanopore Technologies. Briefly, high molecular weight DNA was sheared with a g-TUBE (Covaris) to an average fragment length of 20 kbp. The sheared DNA was repaired using the FFPE repair mix according to the manufacturer’s instructions (New England Biolabs, Ipswich, USA). After cleaning up the DNA with an extraction using a ratio of 0.4:1 Ampure XP beads to DNA the DNA ends were polished and an A overhang was added with the the NEBNext End Prep Module and again cleaned up with an extraction using a ratio of 1:1 Ampure XP beads to DNA the DNA prior to ligation. The adaptor and hairpin adapter were ligated using Blunt/TA Ligase Master Mix (New England Biolabs). The final library was prepared by cleaning up the ligation mix using MyOne C1 beads (Invitrogen).

To prepare 2D libraries for R9 sequencing runs we used the SQK-NSK007 kit from Oxford Nanopore Technologies. The procedure to prepare a library with this kit is largely the same as with the SQK-MAP006 kit. 1D library preparation was done with the SQK-RAD001 kit from Oxford Nanopore Technologies. In short, high molecular weight DNA was tagmented with a transposase. The final library was prepared by ligation of the sequencing adapters to the tagmented fragments using the Blunt/TA Ligase Master Mix (New England Biolabs).

Library preparation for R9.4 sequencing runs was done with the SQK-LSK108 and the SQK-RAD002 kits from Oxford Nanopore Technologies. The procedure to prepare libraries using the SQK-RAD002 kit was the same as for the SQK-RAD001 kit. For SQK-LSK108 the procedure was essentially the same as for SQK-NSK007 except that only adapters and no hairpins were ligated to the DNA fragments. As a consequence the final purification step was done using Ampure XP beads instead of MyOne C1 beads. Libraries for R7.3 and R9 flow cells were directly loaded on the flow cells. To load the library on the R9.4 flow cell the DNA fragments were first bound to beads which were then loaded on the flow cell.

The MinKNOW software was used to control the sequencing process and the read files were uploaded to the cloud based Metrichor EPI2ME platform for base calling. Base called reads were downloaded for further processing and assembly.

### Nanopore read alignment

From the base called read files produced by the Metrichor EPI2ME platform sequence files in FASTA format were extracted using the R-package poRe v0.17 [46]. We used BWA-MEM [47] to align nanopore reads to selected seeds, using specific settings for each nanopore chemistry. The built-in *-x ont2d* setting (*-k 14 -W 20 -r 10 -A 1 -B 1 -O 1 -E 1 -L 0*) is too tolerant for newer chemistries. We therefore optimized alignment settings (*-k* and *-W* only) on small subsets to yield the highest recall (number of aligning reads) at the highest precision (number of seeds detected/number of alignments). With all other settings as before, this yielded the following parameters: *-k 14-W 45* (R7.3 2D); *-k 16 -W 50* (R9 1D); *-k 19 -W 60* (R9 2D); *-k 16 -W 60* (R9.4 1D).

### Genome assembly using TULIP

Currently, TULIP consists of two prototype scripts in Perl: *tulipseed.perl* and *tulipbulb.perl* (version 0.4 ‘European eel’). The *tulipseed* script constructs the seed graph based on input SAM files and a set seed length, and outputs a simplified graph and seed arrangements (scaffold models). *tulipbulb* adds seed and long read sequence to the scaffolds, and exports either a complete set of uncorrected scaffolds, or for each scaffold two separate files: the uncorrected sequence, and a FASTA ‘bundle’ consisting of all long reads associated with that scaffold.

For each scaffold, we used the long read bundle and Illumina data to polish it according to ONT guidelines [48]. We first corrected nanopore-derived scaffolds with nanopore data using Racon [19], based on alignments produced by Graphmap version 0.3.0 [49]. Ultimately Racon sequence correction is performed by SPOA [50], which is a partial order alignment algorithm that generates consensus sequences.

Subsequently, we used previously generated Illumina data (trimmed to Phred 30 quality values using Sickle version 1.33 [51]) in a second correction step using Pilon (version 1.21), an integrated software tool for assembly improvement [33, 34]. Pilon uses evidence from the alignment between short-read data and Racon-corrected scaffolds to identify events that are different in the draft genome compared to the support of short-read data.

All genome assembly steps and analyses were performed on a desktop computer equipped with an Intel Xeon E3-1241 3.5 GHz processor, in a virtual machine (Oracle VirtualBox version 4.3.26) running Ubuntu 16.04 LTS with 28 GB RAM and 4 processor threads available. For the final candidate assembly, the TULIP scripts required a maximum of 4.4 GB RAM.

### Genome alignment

Uncorrected scaffolds were aligned against the 2010 scaffolds using nucmer version 3.23 [52], with settings *--maxmatch* and *--minmatch 100*, filtered for optimal correspondence (delta-filter −1), and visualized using mummerplot (with the *--layout* option). The five largest scaffolds were likewise aligned against the 2012 scaffolds, but with settings encouraging longer alignments (*--breaklen 1000* and *--minmatch 25*) and not filtered. The 285 nt seeds were aligned against the 2012 draft scaffolds using BWA-MEM with default settings.

**List of abbreviations**
bp (kbp, Mbp, Gbp): Basepairs (thousands, millions, billions of basepairs)
N50: The length-weighed median fragment length, such that 50% of the fragment length sum is in fragments larger than the N50
*k*-mer: A sequence of length *k*
C-value: The weight of a haploid genome
CPU: Central processing unit
ONT: Oxford Nanopore Technologies
PacBio: Pacific Biosciences

## Declarations

### Ethics approval

Experiments were approved by the animal ethical commission of Leiden University (DEC #13060).

### Availability of data and materials

Submission of the nanopore and Illumina sequencing data to ENA and NCBI is in progress. The Illumina and nanopore sequencing data can temporarily be accessed at https://surfdrive.surf.nl/files/index.php/s/5wOBiWqqyUZV2Yd

The Racon-and Pilon-corrected candidate assembly is available at http://www.zfgenomics.com/sub/eel

The TULIP-scripts are available at https://github.com/Generade-nl/TULIP

## Authors’ contributions

HJJ, SD, F-AW, WS, AK, APP, BP, HPS, GEvdT, RPD and CVH conceived the research. RPD coordinated the project. HJJ and SAJ-R performed sequencing, ML and CVH assembled the genome, HJJ, RPD and CVH analyzed the data. HJJ, ML, RPD and CVH wrote the paper with input from all other authors.

## Acknowledgements

We are grateful to the DUPAN Foundation for making this project possible, as well as to Rosemary Dokos and Oliver Hartwell at Oxford Nanopore for technical support and encouragement.

---

**Fig. S1.**
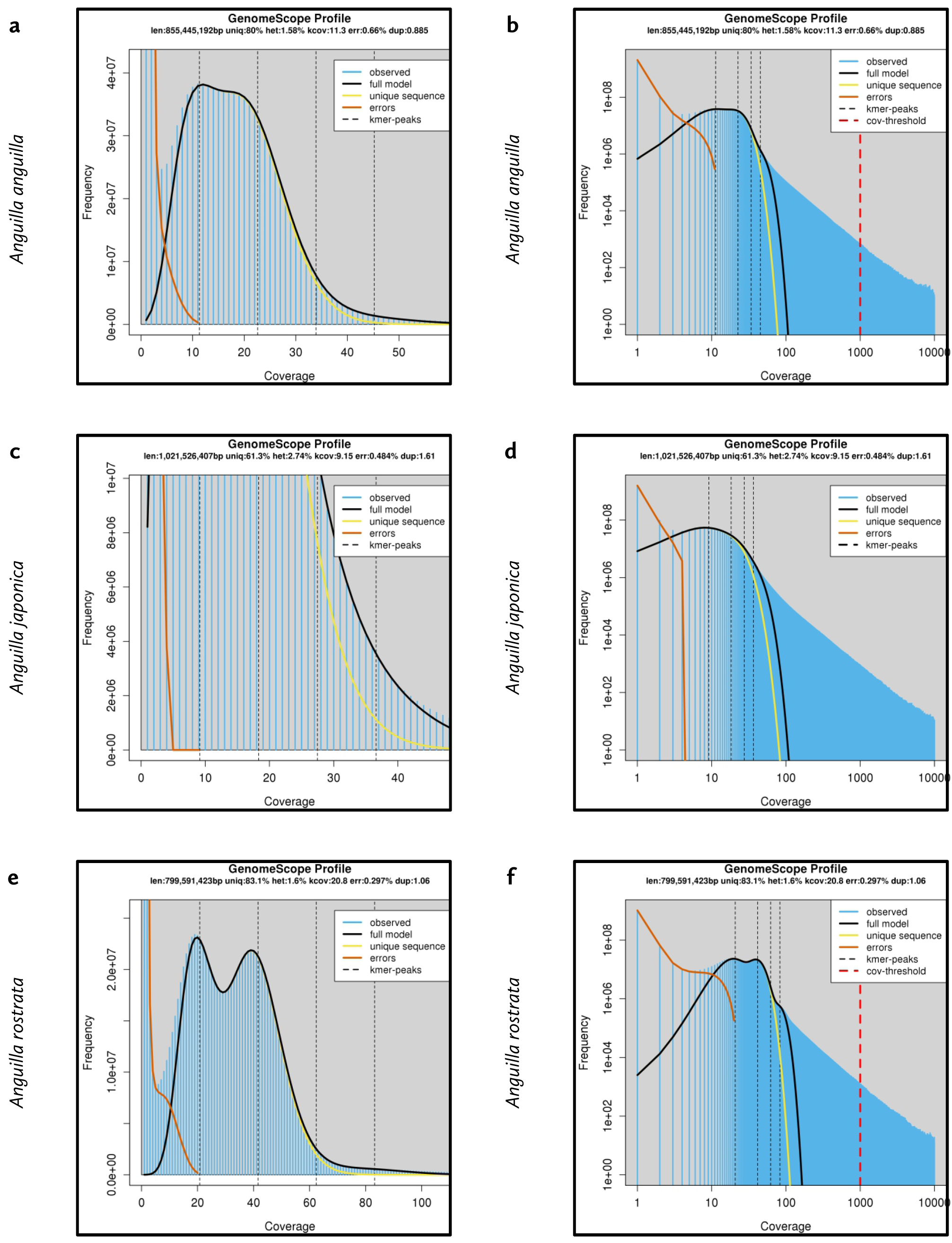
GenomeScope k-mer profiles Shown are the 19-mer profile analyses for **a** A. anguilla, **b** A. japonica and **c** A. rostrata. Both regular and logarithmic scale plots are included. The full analyses are available at the GenomeScope website (http://qb.cshl.edu/genomescope/analysis.php) using the codes TDVyqzdJXugs2lEcd2AB (A. anguilla), VtNZvSlV7nzfq6yvTlAp (A. japonica) and 8citu1cxv9SHXOzqbA43 (A. rostrata).

**Fig. S2.**
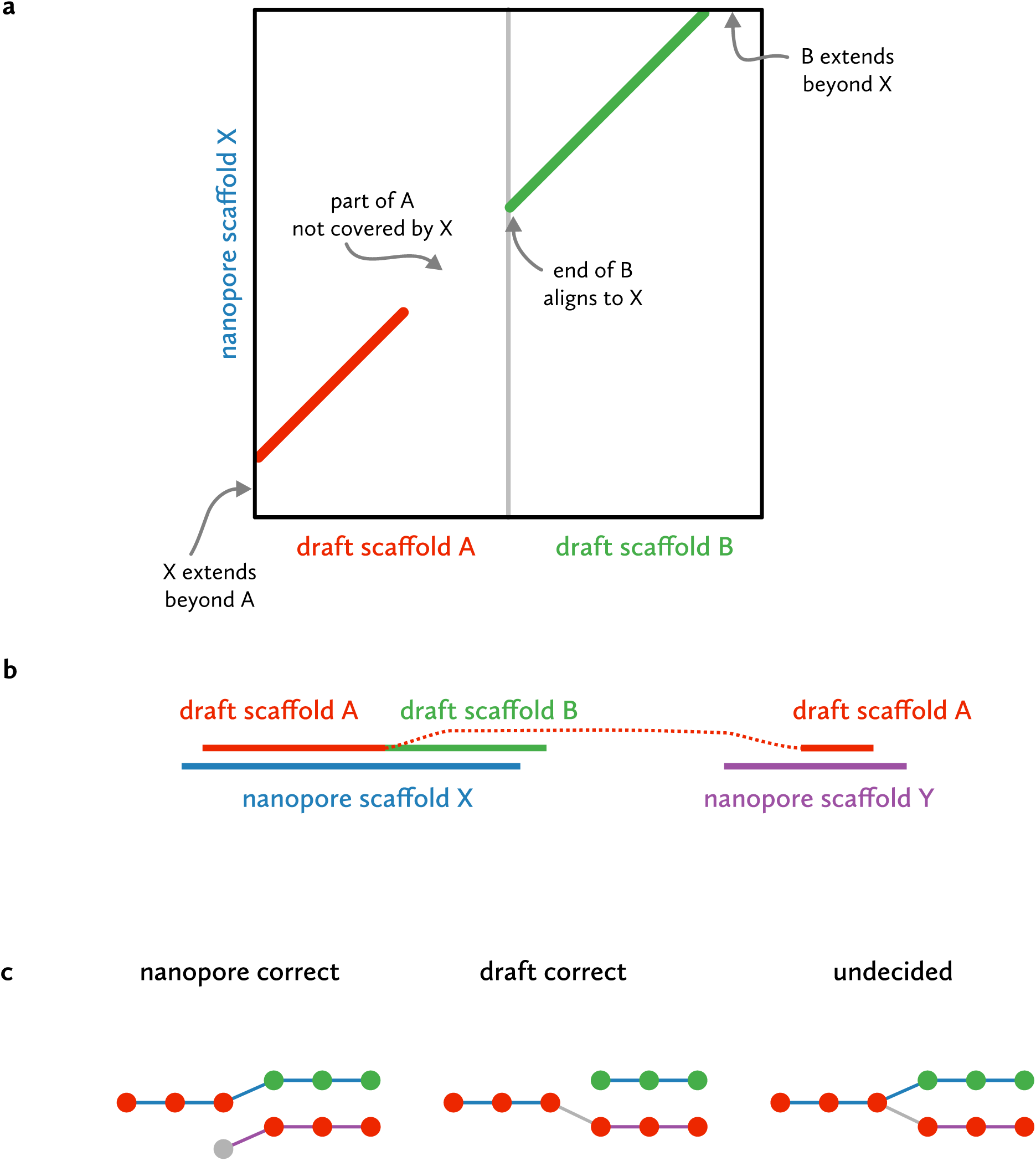
Misassembly scenarios If draft scaffolds do not align completely to a single nanopore scaffold, this is apparent in the alignment plot (**a**). The origins of the actual situation (**b**) can be gleaned from the nanopore graph (**c**). Based on the local graph context around the inconsistency, multiple explanations are possible: nanopore evidence can exist to support the nanopore scaffolds only (in which case the draft scaffold is probably incorrect), to support the draft scaffold only (in which case the nanopore scaffold is incorrect), or to support both (in which case additional evidence needs to be examined to determine the correct scaffolding path).

**Fig. S3.**
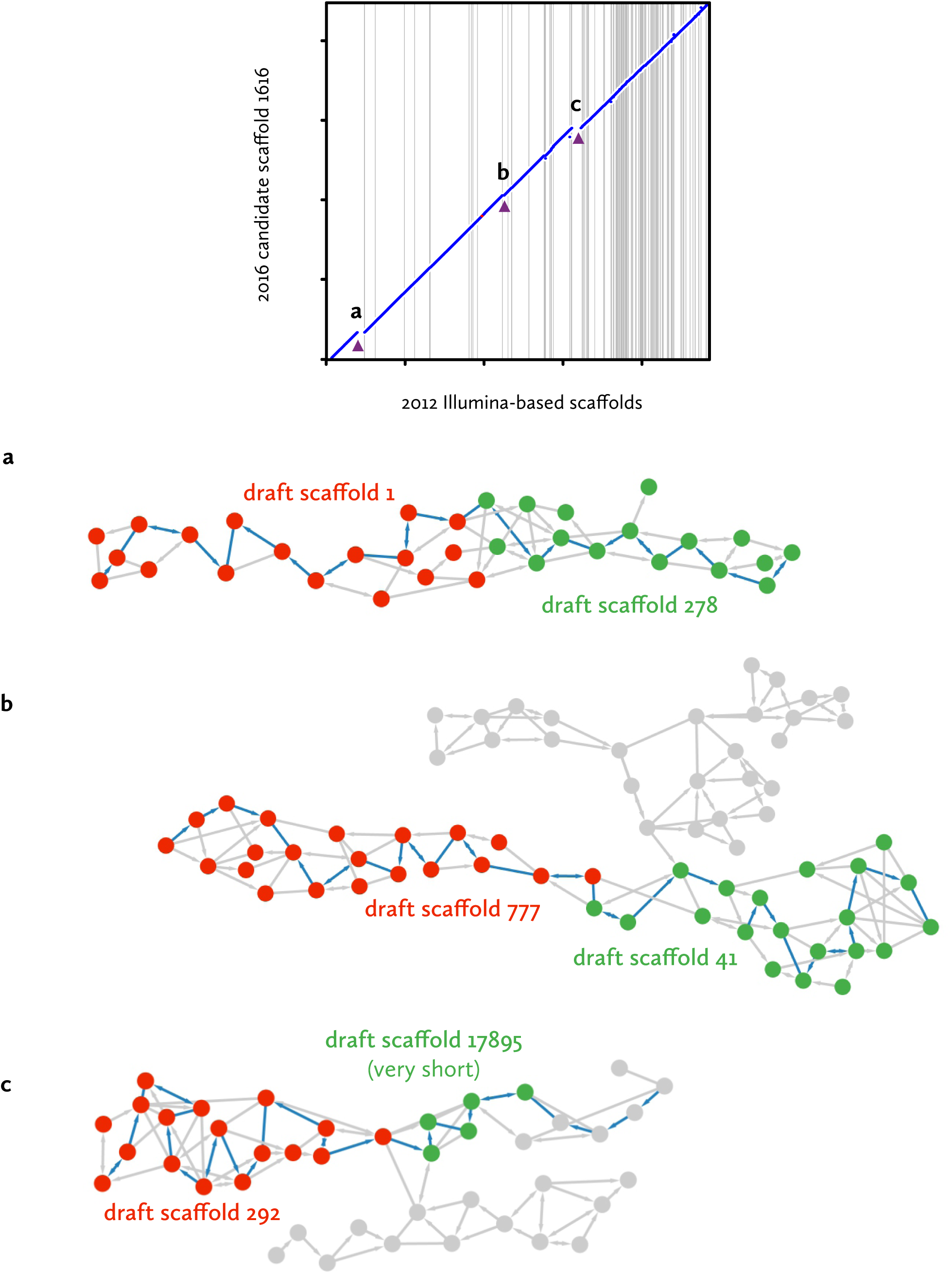
Local graph neighbourhoods of scaffold inconsistencies. For each of the inconsistencies identified in Fig. 5b–f, the local neighbourhood in the initial seed graph is shown (similar to Fig. 3 and Supplementary Fig. 2c). Red and green nodes represent seeds that align to the truncated old scaffold and its non-truncated neighbour, respectively. Grey nodes do not align to these scaffolds (or at least, not locally), yellow nodes align partially to two scaffolds. The final extracted TULIP scaffold paths are indicated by blue arrows. As in the draft the ‘red’ scaffolds do not end at the joins to the ‘green’ scaffolds, an alternative path possibility of continuing with ‘red’ seeds would be expected at this point. In none of the cases examined does this appear to be the case.

**Fig. S4.**
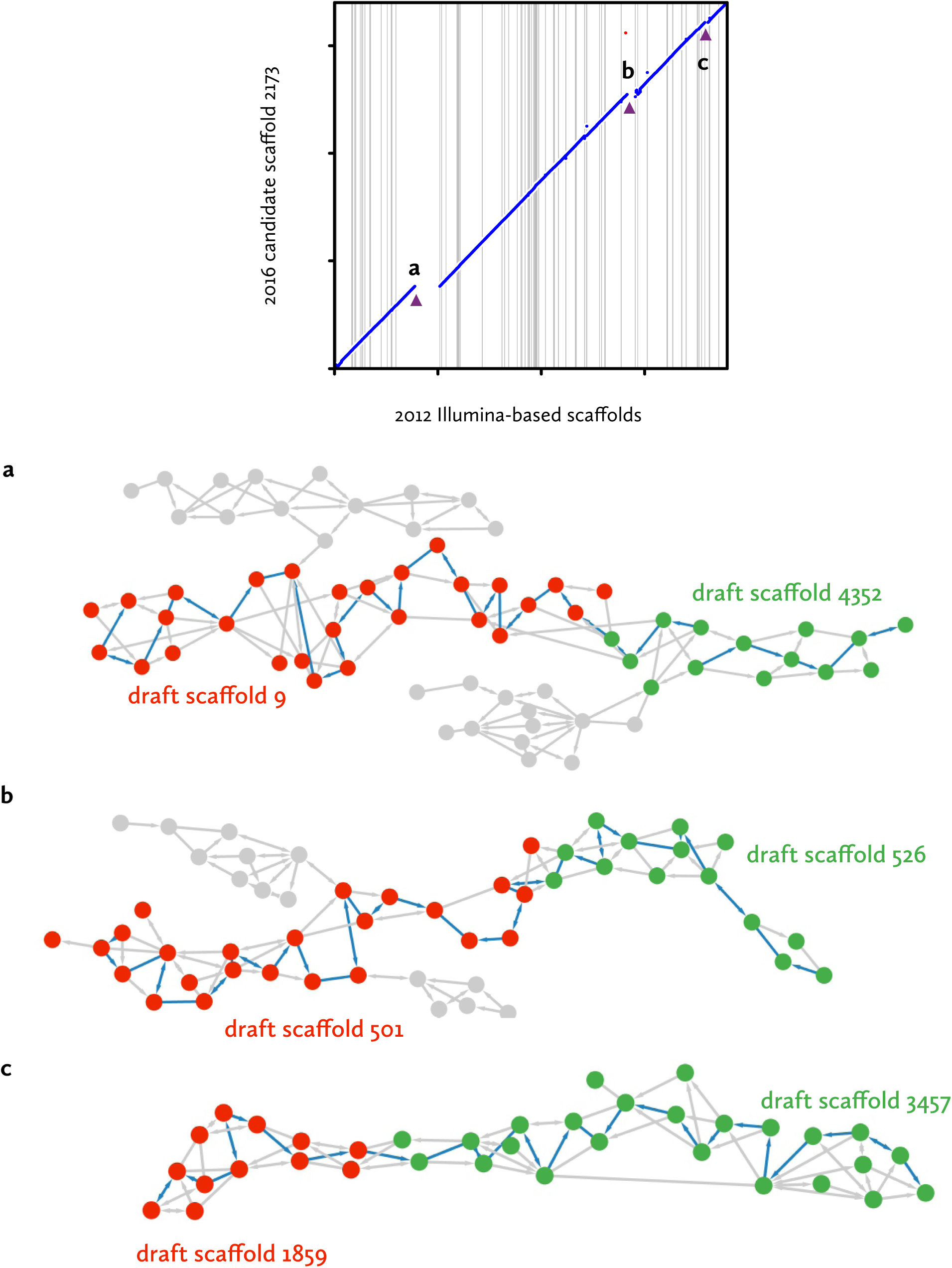
Local graph neighbourhoods of scaffold inconsistencies. For each of the inconsistencies identified in Fig. 5b–f, the local neighbourhood in the initial seed graph is shown (similar to Fig. 3 and Supplementary Fig. 2c). Red and green nodes represent seeds that align to the truncated old scaffold and its non-truncated neighbour, respectively. Grey nodes do not align to these scaffolds (or at least, not locally), yellow nodes align partially to two scaffolds. The final extracted TULIP scaffold paths are indicated by blue arrows. As in the draft the ‘red’ scaffolds do not end at the joins to the ‘green’ scaffolds, an alternative path possibility of continuing with ‘red’ seeds would be expected at this point. In none of the cases examined does this appear to be the case.

**Fig. S5.**
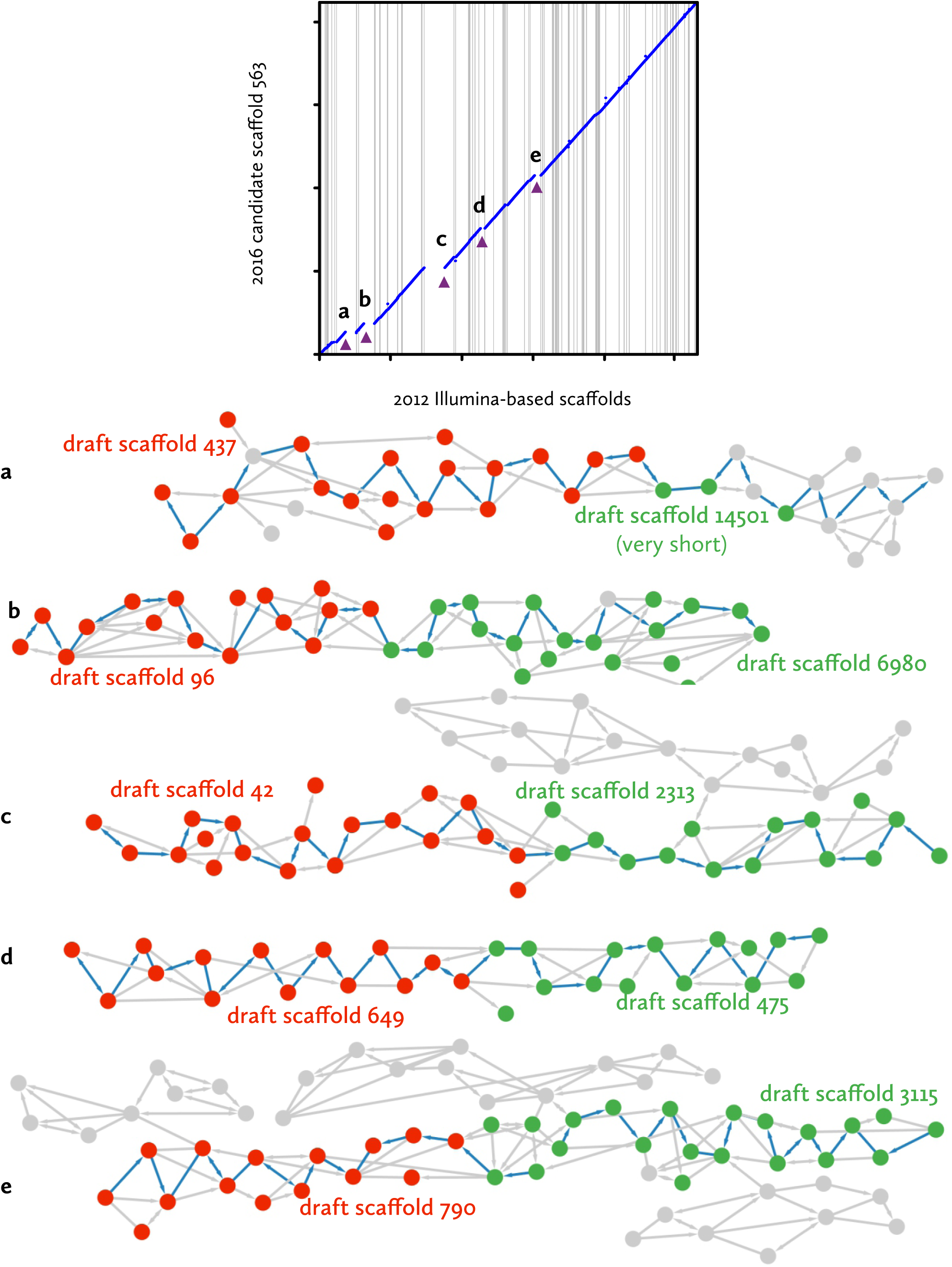
Local graph neighbourhoods of scaffold inconsistencies. For each of the inconsistencies identified in Fig. 5b–f, the local neighbourhood in the initial seed graph is shown (similar to Fig. 3 and Supplementary Fig. 2c). Red and green nodes represent seeds that align to the truncated old scaffold and its non-truncated neighbour, respectively. Grey nodes do not align to these scaffolds (or at least, not locally), yellow nodes align partially to two scaffolds. The final extracted TULIP scaffold paths are indicated by blue arrows. As in the draft the ‘red’ scaffolds do not end at the joins to the ‘green’ scaffolds, an alternative path possibility of continuing with ‘red’ seeds would be expected at this point. In none of the cases examined does this appear to be the case.

**Fig. S6.**
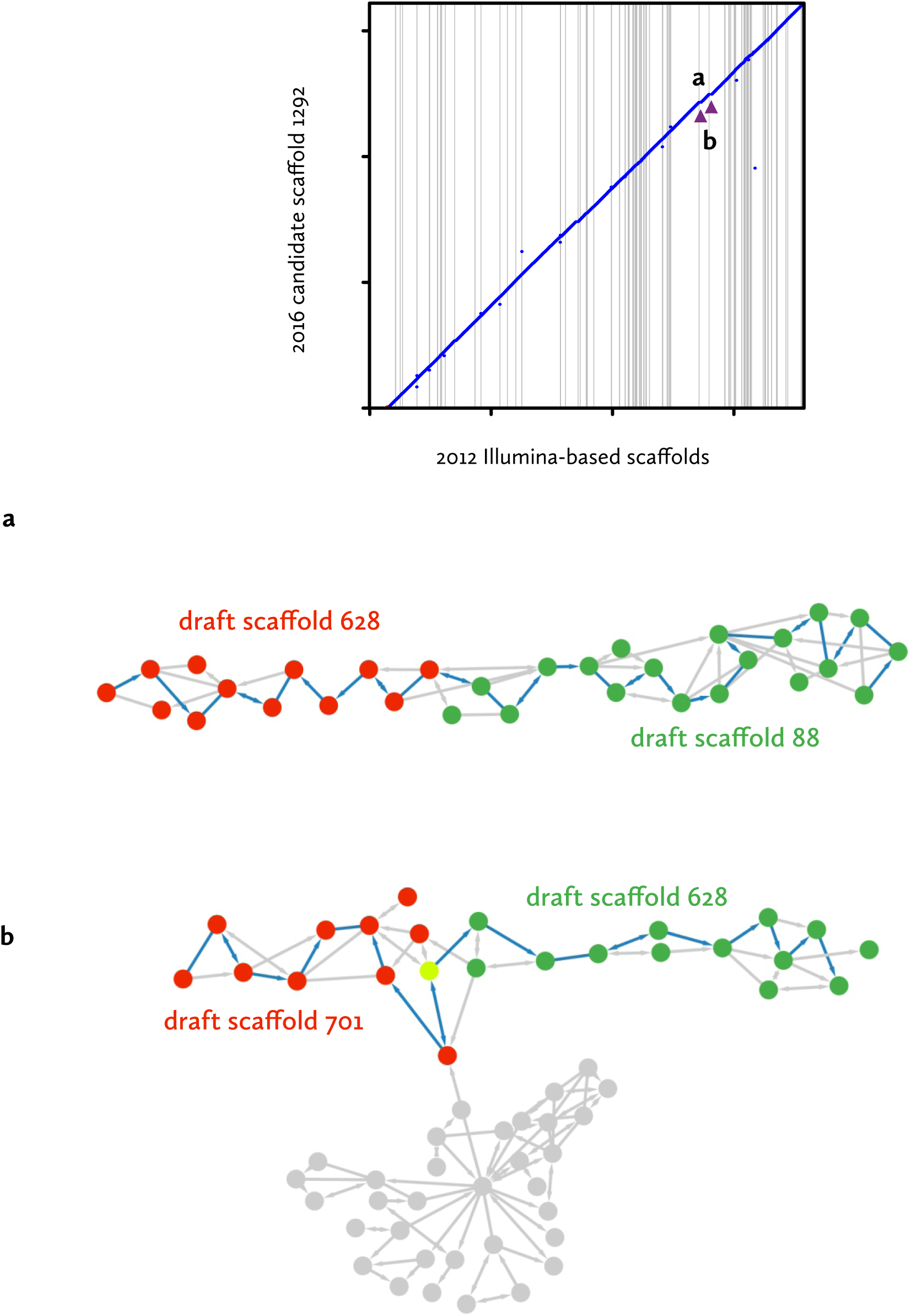
Local graph neighbourhoods of scaffold inconsistencies. For each of the inconsistencies identified in Fig. 5b–f, the local neighbourhood in the initial seed graph is shown (similar to Fig. 3 and Supplementary Fig. 2c). Red and green nodes represent seeds that align to the truncated old scaffold and its non-truncated neighbour, respectively. Grey nodes do not align to these scaffolds (or at least, not locally), yellow nodes align partially to two scaffolds. The final extracted TULIP scaffold paths are indicated by blue arrows. As in the draft the ‘red’ scaffolds do not end at the joins to the ‘green’ scaffolds, an alternative path possibility of continuing with ‘red’ seeds would be expected at this point. In none of the cases examined does this appear to be the case.

**Fig. S7.**
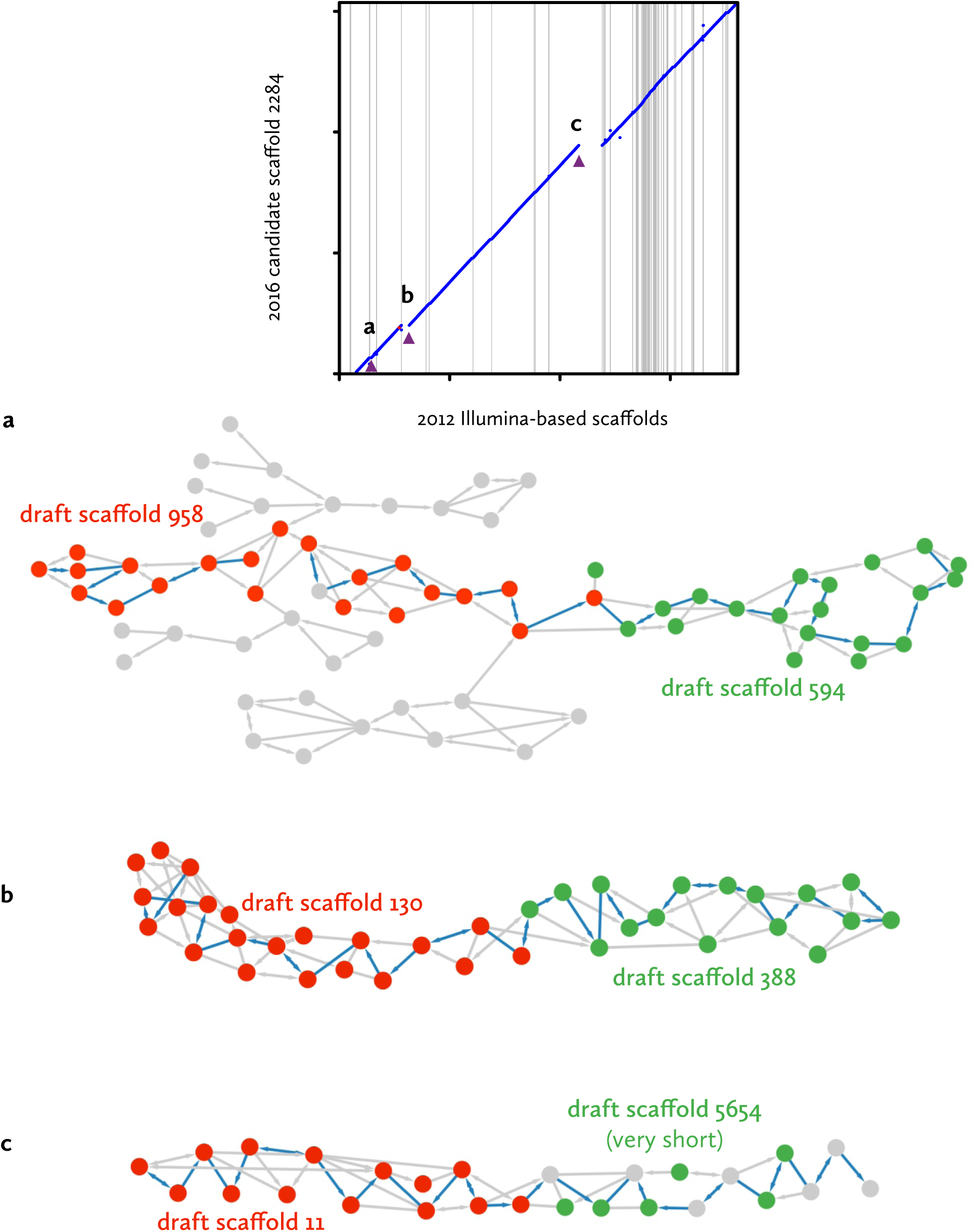
Local graph neighbourhoods of scaffold inconsistencies. For each of the inconsistencies identified in Fig. 5b–f, the local neighbourhood in the initial seed graph is shown (similar to Fig. 3 and Supplementary Fig. 2c). Red and green nodes represent seeds that align to the truncated old scaffold and its non-truncated neighbour, respectively. Grey nodes do not align to these scaffolds (or at least, not locally), yellow nodes align partially to two scaffolds. The final extracted TULIP scaffold paths are indicated by blue arrows. As in the draft the ‘red’ scaffolds do not end at the joins to the ‘green’ scaffolds, an alternative path possibility of continuing with ‘red’ seeds would be expected at this point. In none of the cases examined does this appear to be the case.

## References

1. Coppe A, Pujolar JM, Maes GE, Larsen PF, Hansen MM, Bernatchez L, Zane L, Bortoluzzi S. Sequencing, de novo annotation and analysis of the first Anguilla anguilla transcriptome: EeelBase opens new perspectives for the study of the critically endangered European eel. BMC Genomics. 2010;11:635.

2. Henkel CV, Burgerhout E, de Wijze DL, Dirks RP, Minegishi Y, Jansen HJ, Spaink HP, Dufour S, Weltzien FA, Tsukamoto K, van den Thillart GE. Primitive duplicate Hox clusters in the European eel’s genome. PLoS One. 2012;7:e32231.

3. Pujolar JM, Marino IA, Milan M, Coppe A, Maes GE, Capoccioni F, Ciccotti E, Bervoets L, Covaci A, Belpaire C, Cramb G, Patarnello T, Bargelloni L, Bortoluzzi S, Zane L. Surviving in a toxic world: transcriptomics and gene expression profiling in response to environmental pollution in the critically endangered European eel. BMC Genomics. 2012;13:507.

4. Minegishi Y, Henkel CV, Dirks RP, van den Thillart GE. Genomics in eels – towards aquaculture and biology. Mar Biotechnol (NY). 2012;14:583–90.

5. IUCN Red List. 2014; doi:10.2305/IUCN.UK.2014-1.RLTS.T60344A45833138.en

6. Ager-Wick E, Dirks RP, Burgerhout E, Nourizadeh-Lillabadi R, de Wijze DL, Spaink HP, van den Thillart GE. Tsukamoto K Dufour S, Weltzien FA, Henkel CV. The pituitary gland of the European eel reveals massive expression of genes involved in the melanocortin system. PLoS One. 2013;8:e77396.

7. Dirks RP, Burgerhout E, Brittijn SA, de Wijze DL, Ozupek H, Tuinhof-Koelma N, Minegishi Y, Jong-Raadsen SA, Spaink HP, van den Thillart GE. Identification of molecular markers in pectoral fin to predict artificial maturation of female European eels (Anguilla anguilla). Gen Comp Endocrinol. 2014;204:267–76.

8. Burgerhout E, Minegishi Y, Brittijn SA, de Wijze DL, Henkel CV, Jansen HJ, Spaink HP, Dirks RP, van den Thillart GE. Changes in ovarian gene expression profiles and plasma hormone levels in maturing European eel (Anguilla anguilla); biomarkers for broodstock selection. Gen Comp Endocrinol. 2016;225:185–96.

9. Churcher, AM, Hubbard, PC, Marques, JP, Canário, AV, Huertas, M. Deep sequencing of the olfactory epithelium reveals specific chemosensory receptors are expressed at sexual maturity in the European eel Anguilla anguilla. Mol Ecol. 2015; 24:822–34.

10. Pelster, B, Schneebauer, G, Dirks, RP. Anguillicola crassus infection significantly affects the silvering related modifications in steady state mRNA levels in gas gland tissue of the European eel. Front Physiol. 2016;7:175.

11. Pujolar JM, Jacobsen MW, Frydenberg J, Als TD, Larsen PF, Maes GE, Zane L, Jian JB, Chench L, Hansen MM. A resource of genome-wide single-nucleotide polymorphisms generated by RAD tag sequencing in the critically endangered European eel. Mol Ecol Resour. 2013;13:706–14.

12. Pasquier J, Lafont AG, Jeng SR, Morini M, Dirks R, van den Thillart G, Tomkiewicz J, Tostivint H, Chang CF, Rousseau K, Dufour S. Multiple kisspeptin receptors in early osteichthyans provide new insights into the evolution of this receptor family. PLoS One. 2012;7:e48931.

13. Maugars G, Dufour S. Demonstration of the coexistence of duplicated LH receptors in teleosts, and their origin in ancestral actinopterygians. PLoS One. 2015;10:e0135184.

14. Morini M, Pasquier J, Dirks R, van den Thillart G, Tomkiewicz J, Rousseau K, Dufour S, Lafont AG. Duplicated leptin receptors in two species of eel bring new insights into the evolution of the leptin system in vertebrates. PLoS One. 2015;10:e0126008.

15. Pujolar JM, Jacobsen MW, Als TD, Frydenberg J, Munch K, Jónsson B, Jian JB, Chench L, Maes GE, Bernatchez L, Hansen MM. Genome-wide single-generation signatures of local selection in the panmictic European eel. Mol Ecol. 2014;23:2514–28.

16. Jacobsen MW, Pujolar JW, Bernatchez L, Munch K, Jian J, Niu Y, Hansen MM. Genomic footprints of speciation in Atlantic eels (Anguilla anguilla and A. rostrata). Mol Ecol. 2014;23:4785–4798.

17. Ip CL, Loose M, Tyson JR, de Cesare M, Brown BL, Jain M, Leggett RM, Eccles DA, Zalunin V, Urban JM, Piazza P, Bowden RJ, Paten B, Mwaigwisya S, Batty EM, Simpson JT, Snutch TP, Birney E, Buck D, Goodwin S, Jansen HJ, O'Grady J, Olsen HE, Min ION, Analysis and Reference Consortium. MinION Analysis and Reference Consortium: Phase 1 data release and analysis. F1000Res.2015;4:1075.

18. Jain M, Olsen HE, Paten B, Akeson M. The Oxford Nanopore MinION: delivery of nanopore sequencing to the genomics community. Genome Biol. 2016;17:239.

19. Vaser R, SovićI, Nagarajan N, Šikić M. Fast and accurate de novo genome assembly from long uncorrected reads.BioRxiv. 2016; doi:10.1101/068122.

20. Koren S, Harhay GP, Smith TP, Bono JL, Harhay DM, Mcvey SD, Radune D, Bergman NH, Phillippy AM. Reducing assembly complexity of microbial genomes with single-molecule sequencing. Genome Biol. 2013;14:R101

21. Loman NJ, Quick J, Simpson JT. A complete bacterial genome assembled de novo using only nanopore sequencing data. Nat Methods. 2015;12:733–5.

22. Koren S, Walenz BP, Berlin K, Miller JR, Phillippy AM. Canu: scalable and accurate long-read assembly via adaptive k-mer weighting and repeat separation.BioRxiv. 2016; doi:10.1101/071282.

23. Myers, G. https://dazzlerblog.wordpress.com. Accessed December 2016.

24. Chin CS, Peluso P, Sedlazeck FJ, Nattestad M, Concepcion GT, Clum A, Dunn C, O’Malley R, Figueroa-Balderas R, Morales-Cruz A, Cramer GR, Delledonne M, Luo C, Ecker JR, Cantu D, Rank DR, Schatz MC. Phased diploid genome assembly with single-molecule real-time sequencing. Nat Methods. 2016;13:1050–1054.

25. Li H. Minimap and miniasm: fast mapping and de novo assembly for noisy long sequences. Bioinformatics. 2016;32:2103–10.

26. Kamath GM, Shomorony I, Xia F, Courtade TA, Tse DN. HINGE: long-read assembly achieves optimal repeat resolution. BioRxiv. 2016; doi:10.1101/062117

27. Keinath MC, Timoshevskiy VA, Timoshevskaya NY, Tsonis PA, Voss SR, Smith JJ. Initial characterization of the large genome of the salamander Ambystoma mexicanum using shotgun and laser capture chromosome sequencing. Sci Rep. 2015;5:16413.

28. Biscotti MA, Gerdol M, Canapa A, Forconi M, Olmo E, Pallavicini A, Barucca M, Schartl M. The lungfish transcriptome: a glimpse into molecular evolution events at the transition from water to land. Sci Rep. 2016;6:21571.

29. Zonneveld BJ. The systematic value of nuclear genome size for all species of Tulipa L. (Liliacaeae). Plant Syst Evol. 2009; 281:217–45.

30. Gregory, TR. Animal genome size database. http://www.genomesize.com. Accessed November 2016.

31. Li X, Waterman MS. Estimating the repeat structure and length of DNA sequences using l-tuples. Genome Res. 2003;13;1916–22.

32. Vuture GW, Sedlazeck FJ, Nattestad M, Underwood CJ, Fang H, Gurtowki J, Schatz MC. GenomeScope: fast reference-free genome profiling from short reads. BioRxiv. 2016;doi:10.1101/075978.

33. Walker BJ, Abeel T, Shea T, Priest M, Abouelliel A, Sakthikumar S, Cuomo CA, Zeng Q, Wortman J, Young SK, Earl AM. Pilon: an integrated tool for comprehensive microbial variant detection and genome assembly improvement. PLoS One. 2014;9:e112963.

34. Goodwin S, Gurtowski J, Ethe-Sayers S, Deshpande P, Schatz MC, McCombie WR. Oxford Nanopore sequencing, hybrid error correction, and de novo assembly of a eukaryotic genome. Genome Res. 2015;25;1–7.

35. Boetzer M, Henkel CV, Jansen HJ, Butler D, Pirovano W Scaffolding pre-assembled contigs using SSPACE. Bioinformatics. 2011;27:578–9.

36. Henkel CV,Dirks RP, de Wijze DL, Minegishi Y, Aoyama J, Jansen HJ, Turner B, Knudsen B, Bundgaard M, Hvam KL, Boetzer M, Pirovano W, Weltzien FA, Dufour S, Tsukamoto K, Spaink HP, van den Thillart GE. First draft genome sequence of the Japanese eel, Anguilla japonica. Gene. 2012;511:195–201.

37. Pavey SA, Laporte M, Normandeau E, Gaudin J, Letourneau L, Boisvert S, Corbeil J, Audet C, Bernatchez L. Draft genome of the American eel (Anguilla rostrata). Mol Ecol Resour. 2016;doi:10.1111/1755-0998.12608.

38. Albert, V, Jónsson, B, Bernatchez, L. Natural hybrids in Atlantic eels (Anguilla anguilla, A. rostrata): evidence for successful reproduction and fluctuating abundance in space and time. Mol Ecol. 2006;15:1903–16.

39. Hunt M, Newbold C, Berriman M, Otto TD. A comprehensive evaluation of assembly scaffolding tools. Genome Biol. 2014;15:R42.

40. Chapman JA, Maschner M, Buluç A, Barry K, Georganas E, Session A, Strnadova V, Jenkins J, Sehgal S, Oliker L, Schmutz J, Yelick KA, Scholz U, Waugh R, Poland JA, Muehlbauer GJ, Stein N, Rokhsar D. A whole-genome shotgun approach for assembling and anchoring the hexaploidy bread wheat genome. Genome Biol. 2015;16:26.

41. Urban JM, Bliss J, Lawrence CE, Gerbi SA., Sequencing ultra-long DNA, molecules with the Oxford Nanopore MinION. BioRxiv. 2015; doi:10.1101/019281.

42. Datema E, Hulzink RJ, Blommer L, Valle-Inclan JE, van Orsouw N, Wittenberg AH, de Vos M. The megabase-sized fungal genome of Rhizoctonia solani assembled from nanopore reads only. BioRxiv. 2016; doi:10.1101/084772.

43. Mostovoy Y, Levy-Sakin M, Lam J, Lam ET, Hastie AR, Marks P, Lee J, Chu C, Lin C, Džakula Ž, Cao H, Schlebusch SA, Giorda K, Schnall-Levin M, Wall JD, Kwok PY. A hybrid approach for de novo human genome sequence assembly and phasing. Nat Methods. 2016;13:587–90.

44. Marçais G, Kingsford C. A fast, lock-free approach for efficient parallel counting of occurrences of k-mers. Bioinformatics. 2011; 27:764–70.

45. Magoc T, Salzberg S. FLASH: Fast length adjustment of short reads to improve genome assemblies. Bioinformatics. 2011;27:2957–63.

46. Watson M, Thomson M, Risse J, Talbot R, Santoyo-Lopez J, Gharbi K, Blaxter M. poRe: an R package for the visualization and analysis of nanopore sequencing data. Bioinformatics. 2015;31:114–5.

47. Li H, Durbin R. Fast and accurate long-read alignment with Burrows-Wheeler transform. Bioinformatics. 2010;26:589–95.

48. Oxford Nanopore Technologies.Hybrid assembly pipeline.https://github.com/nanoporetech/ont-assembly-polish. Accessed December 2016.

49. Sović I, Šikić M, Wilm A, Fenlon SN, Chen S, Nagarajan N. Fast and sensitive mapping of nanopore sequencing reads with GraphMap. Nat Commun. 2016;7:11307.

50. Lee, C. Generating consensus sequences from partial order multiple sequence alignment graphs. Bioinformatics. 2003;19:999–1008.

51. Joshi NA, Fass JN. Sickle: A sliding-window, adaptive, quality-based trimming tool for FastQ files. https://github.com/najoshi/sickle. Accessed December 2016.

52. Kurtz S, Phillippy A, Delcher AL, Smoot M, Shumway M, Antonescu C, Salzberg SL. Versatile and open software for comparing large genomes. Genome Biol. 2004;5:R12.

